# CD4+ and CD8+ T cells are required to prevent SARS-CoV-2 persistence in the nasal compartment

**DOI:** 10.1101/2024.01.23.576505

**Authors:** Meenakshi Kar, Katherine E.E. Johnson, Abigail Vanderheiden, Elizabeth J. Elrod, Katharine Floyd, Elizabeth Geerling, E. Taylor Stone, Eduardo Salinas, Stephanie Banakis, Wei Wang, Shruti Sathish, Swathi Shrihari, Meredith E. Davis-Gardner, Jacob Kohlmeier, Amelia Pinto, Robyn Klein, Arash Grakoui, Elodie Ghedin, Mehul S. Suthar

## Abstract

SARS-CoV-2 is the causative agent of COVID-19 and continues to pose a significant public health threat throughout the world. Following SARS-CoV-2 infection, virus-specific CD4+ and CD8+ T cells are rapidly generated to form effector and memory cells and persist in the blood for several months. However, the contribution of T cells in controlling SARS-CoV-2 infection within the respiratory tract are not well understood. Using C57BL/6 mice infected with a naturally occurring SARS-CoV-2 variant (B.1.351), we evaluated the role of T cells in the upper and lower respiratory tract. Following infection, SARS-CoV-2-specific CD4+ and CD8+ T cells are recruited to the respiratory tract and a vast proportion secrete the cytotoxic molecule Granzyme B. Using antibodies to deplete T cells prior to infection, we found that CD4+ and CD8+ T cells play distinct roles in the upper and lower respiratory tract. In the lungs, T cells play a minimal role in viral control with viral clearance occurring in the absence of both CD4+ and CD8+ T cells through 28 days post-infection. In the nasal compartment, depletion of both CD4+ and CD8+ T cells, but not individually, results in persistent and culturable virus replicating in the nasal compartment through 28 days post-infection. Using *in situ* hybridization, we found that SARS-CoV-2 infection persisted in the nasal epithelial layer of tandem CD4+ and CD8+ T cell-depleted mice. Sequence analysis of virus isolates from persistently infected mice revealed mutations spanning across the genome, including a deletion in ORF6. Overall, our findings highlight the importance of T cells in controlling virus replication within the respiratory tract during SARS-CoV-2 infection.

## MAIN

The global impact of Severe Acute Respiratory Syndrome Coronavirus 2 remains devastating, with over 773 million confirmed cases and around 7 million deaths reported worldwide by the end of December 2023 (World Health Organization COVID-19 dashboard; https://covid19.who.int). SARS-CoV-2 is primarily transmitted through respiratory droplets and targets ciliated epithelial cells in the nasal cavity, trachea, and lungs^1,2^. SARS-CoV-2 primarily infects epithelial cells within the respiratory tract by utilizing the cellular receptor angiotensin-converting enzyme 2 (ACE2)^3–5^. Infection of the upper respiratory tract is generally associated with a milder disease outcome whereas dissemination to the lungs, in particular infection of the bronchi, bronchioles, and alveoli, can cause pneumonia, severe disease, acute respiratory distress syndrome and death^3^.

The development of mouse models of SARS-CoV-2 has enabled the study of transmission, immunity, and pathogenesis of this virus^6–10^. The ancestral SARS-CoV-2 strain does not replicate in conventional laboratory mice due to inefficient spike protein binding to the murine ACE2^11,12^. To overcome this limitation, several mouse models have been developed, including human ACE2 (hACE2) transgenic mice that express hACE2 transiently after transduction of hACE2 with viral vectors (*e.g*., Adenovirus)^13,14^, K18-hACE2 transgenic mice^15^), or the use of mouse-adapted strains^7,16,17^. A naturally occurring spike mutation at position N501Y, which is found in many SARS-CoV-2 variants (Alpha, Beta, Gamma and Mu variants), increases the affinity of SARS-CoV-2 spike protein for the murine ACE2 receptor and allows for infection of inbred mice^11^. Infection of conventional laboratory mice with naturally occurring N501Y spike mutations show inflammatory infiltrates, alveolar edema, and alveolitis^13,16,18,19^. Using this model, we recently identified that the CCR2-monocyte signaling axis is important for controlling virus replication and dissemination within the lungs and protection against the SARS-CoV-2 B.1.351 (“Beta”) variant^18^.

Following SARS-CoV-2 infection, both CD4+ and CD8+ T cells are detectable within the peripheral blood of patients with COVID-19^20–23^. These circulating virus-specific CD4+ and CD8+ T cells target several SARS-CoV-2 proteins, including Spike and Nucleocapsid, and are polyfunctional and durable with estimated half-lives of over 200 days^20^. SARS-CoV-2-specific CD4+ T cells are important in promoting antibody responses and mitigating disease severity as reduced responses associate with increased disease severity^24–27^. CD8+ T cells appear to play a protective role with reduced responses correlating with adverse prognoses^28^. Rapid type I interferon (IFN) responses and virus-specific CD8+ T cell responses coincide with milder SARS-CoV-2 infections, preceding the development of antibodies by one to two weeks^29^. However, the role of CD4+ and CD8+ T cells within the respiratory tract is not well understood and is only beginning to be studied in the context of SARS-CoV-2 infection in humans^30,31^.

In this study, we investigated the role of CD4+ and CD8+ T cells on SARS-CoV-2 infection in both the upper respiratory tract (i.e., nasal turbinates) and the lower respiratory tract. Following infection, the respiratory tract recruits SARS-CoV-2-specific CD4+ and CD8+ T cells, a significant proportion of which release the cytotoxic substance Granzyme B. Through T cell depletion using antibodies prior to infection, we discovered that CD4+ and CD8+ T cells have distinct roles in the upper and lower respiratory tract. In the lungs, T cells have a limited impact on viral control, as viral clearance occurs even in the absence of both CD4+ and CD8+ T cells up to 28 days post-infection (pi). Conversely, in the nasal compartment, depleting both CD4+ and CD8+ T cells (but not individually) leads to persistent and culturable virus replication in the nasal compartment for 28 days pi. Utilizing *in situ* hybridization, we observed persistent SARS-CoV-2 infection in the nasal epithelial layer of mice depleted of both CD4+ and CD8+ T cells. Sequence analysis of virus isolates from persistently infected mice revealed mutations spanning the entire genome. In summary, our findings underscore the crucial role of T cells in controlling virus replication within the respiratory tract during SARS-CoV-2 infection.

## RESULTS

### SARS-CoV-2 infection kinetics and immune response in C57BL/6 mice

The emergence of SARS-CoV-2 variants encoding an N501Y mutation within the spike protein allows for productive infection of conventional laboratory mice. Previously, we had shown that infection of C57BL/6 mice with the B.1.351 strain results in virus replication in the lungs that corresponds with induction of innate immune responses^18,32^. To further characterize this model, we infected C57BL/6 mice with doses of B.1.351 ranging from 10^5^ to 10^6^ PFU through the intranasal route. We found that increasing virus inoculum led to increased weight loss with mice infected with 5 x 10^6^ PFU showing nearly 20% weight loss with 30% mortality (**Fig. 1a**). For subsequent experiments, we selected 1 x 10^6^ PFU as the viral dose as we observed consistent weight loss in the absence of mortality. We next evaluated the kinetics of virus replication in the upper and lower respiratory tracts. Infectious virus in the lungs, as measured by plaque assay on Vero-ACE2-TMPRSS2 over-expressing cells, peaked between days 2 and 4 post-infection (pi) and reached undetectable levels by day 10 pi (**Fig. 1b**). As a more sensitive measure of virus replication, we measured viral RNA-dependent RNA polymerase (RdRp) levels by qRT-PCR and observed 116-fold reduction in virus replication in the lungs and nasal turbinates (**Fig. 1c**). A hallmark of SARS-CoV-2 infection is the generation of systemic spike- and nucleocapsid-specific antibody responses^33^. Following infection, B.1.351-infected mice generated robust IgG responses against the receptor-binding domain (RBD) (GMT 107839), Spike (GMT 249211) and Nucleocapsid (GMT 39803) that corresponded to live virus neutralization activity (FRNT_50_ GMT 175) (**Fig. 1d-f**).

**Figure 1.**
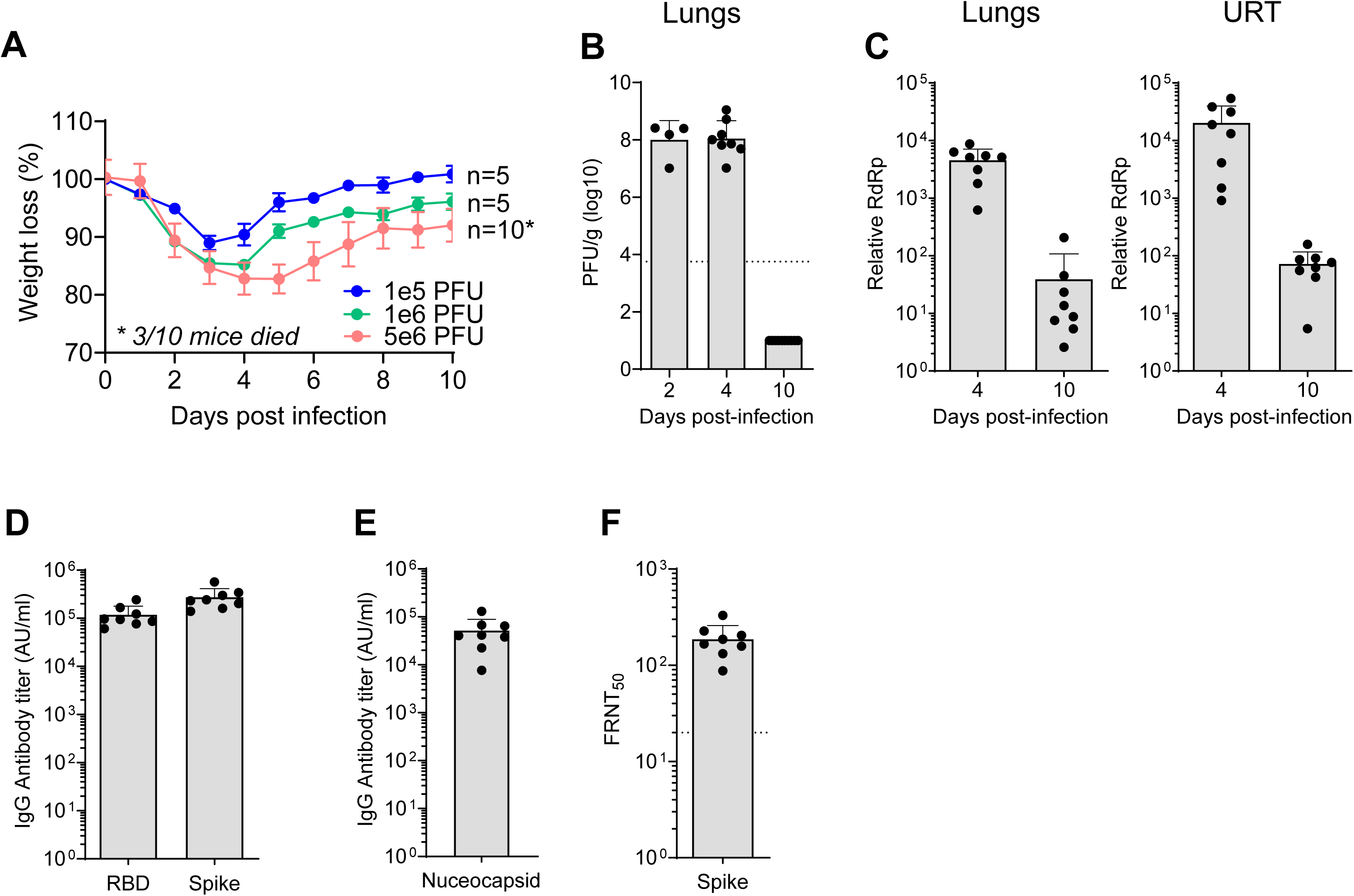
SARS-CoV-2 infection kinetics and immune response in C57BL/6 mice. C57BL/6 mice were infected with the SARS-CoV-2 B.1.351 (Beta) variant or an equal volume of saline for mock mice (A) Percent of initial weight for Beta infected mice at indicated PFUs over ten days. (B) Quantification of infectious virus at indicated days post-infection as measured by plaque assay and expressed as PFU/gm of lung tissue. (C) Quantification of viral RNA by qRT-PCR for SARS-CoV-2 RNA-dependent RNA polymerase (RdRp) in lungs (left) and nasal turbinates (right). CT values represented as relative fold change over mock (log10). IgG antibody titers against (D) RBD and spike and (E) nucleocapsid as measured by an electrochemiluminescent multiplex immunoassay and reported as arbitrary units per ml (AU/mL) and normalized by a standard curve for the B.1.351 SARS-CoV-2 variant. (F) Neutralizing antibody response measured as 50% inhibitory titer (FRNT50) by focus reduction neutralization assay. Graphs show mean ± SD. Results are representative of data from two independent experiments. Day 2 pi (n=4), day 4 pi (n=8), day 10 pi (n=10).

### SARS-CoV-2 B.1.351 infection triggers antigen-specific T cell responses in the upper and lower respiratory tract

Previous studies in humans have shown that SARS-CoV-2 infection can trigger antigen-specific T cell responses within the nasal compartment, lungs, and periphery (e.g. blood)^20,30,34^. We next evaluated T cell responses within the respiratory tract and periphery of SARS-CoV-2 infected mice. Following B.1.351 infection, we isolated immune cells from the nasal compartment, lungs, and spleen followed by *ex vivo* peptide restimulation to identify antigen-specific CD4+ and CD8+ T cells. Prior to harvesting tissues, mice were intravitally labelled with CD45 antibody conjugated to phycoerythrin (PE) to allow identification of circulating (CD45-PE positive) and tissue-resident/ parenchymal cells (CD45-PE negative) within the respiratory tract (gating strategy shown in **Extended Data Fig. 1**). On day 7 pi, within the spleen, we observed a modest, yet significant, increase in the frequency of CD8+ T cells, but not CD4+ T cells (**Fig. 2a**). However, we observed a significant increase in the frequency of tissue-resident CD8+ T cells, but not CD4+ T cells, within the lungs and nasal compartment (**Fig. 2b-c**). This corresponded to an increase in the total number of CD8+ T cells by 32.3-fold within the nasal compartment and by 3.4-fold in the lungs.

**Figure 2.**
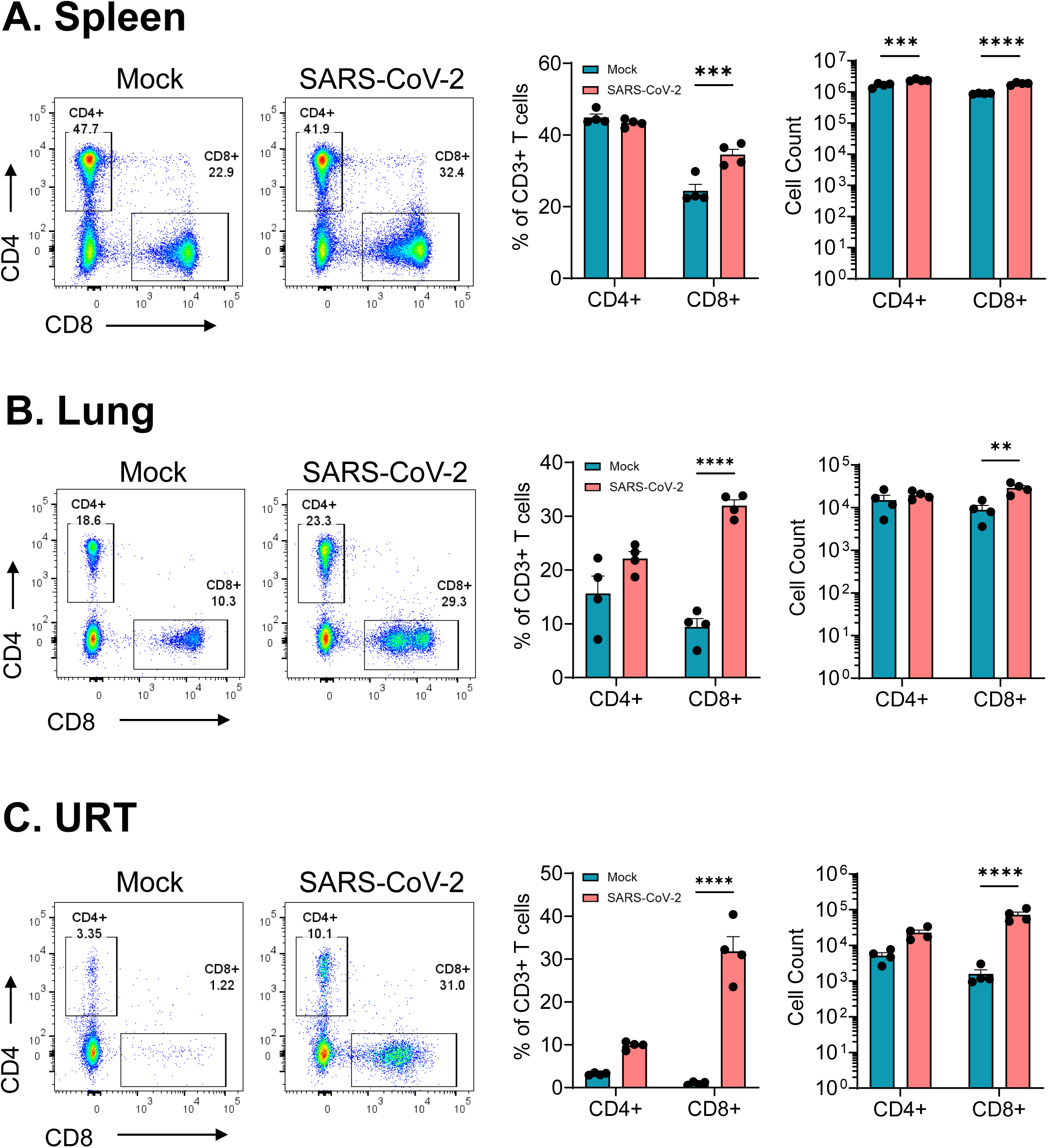
SARS-CoV-2 B.1.351 infection leads to increased infiltration of CD8+ T cells in the respiratory tract but not in the periphery. C57BL/6 mice were infected with the SARS-CoV-2 B.1.351 (Beta) variant at 10^6^ PFU intranasally and at day 7 pi spleen, lungs and URT tissues were harvested, processed for flow cytometry and analyzed via FlowJo. Frequency and cell count for CD4+ and CD8+ T cells in (A) Spleen, (B) Lungs, and (C) URT; representative flow plots on the left, frequency of cells in the middle and cell counts on the right. Graphs show mean ± SD. A two-way ANOVA statistical test was performed. *P < 0.05; **P < 0.01; ***P < 0.001; ****P < 0.0001; no symbol, not significant. Results are representative of data from three independent experiments with 5 mice per group.

We next evaluated the antigen-specific T cell responses by performing *ex vivo* peptide restimulation with a spike peptide pool followed by intracellular staining (**Fig. 3a**). For antigen-specific CD4+ T cells, in both the lungs and nasal compartment, we observed higher cell frequencies and counts of Granzyme B secreting as compared to cytokine secreting cells (**Fig. 3b**). Specifically, we observed 2.35% GrB+, 0.3% TNFα+, 0.67%IFNγ+, and 0.24% TNFα+IFNγ+ CD4+ T cells in the lungs and 6.45% GrB+, 0.56% TNFα+, 0.56%IFNγ+, and 0% TNFα+IFNγ+ CD4+ T cells in the nasal compartment. In the spleen, we observed 0.27% GrB+, 0.50% TNFα+, 0.57%IFNγ+, and 0.08%TNFα+IFNγ+ CD4+ T cells.

**Figure 3.**
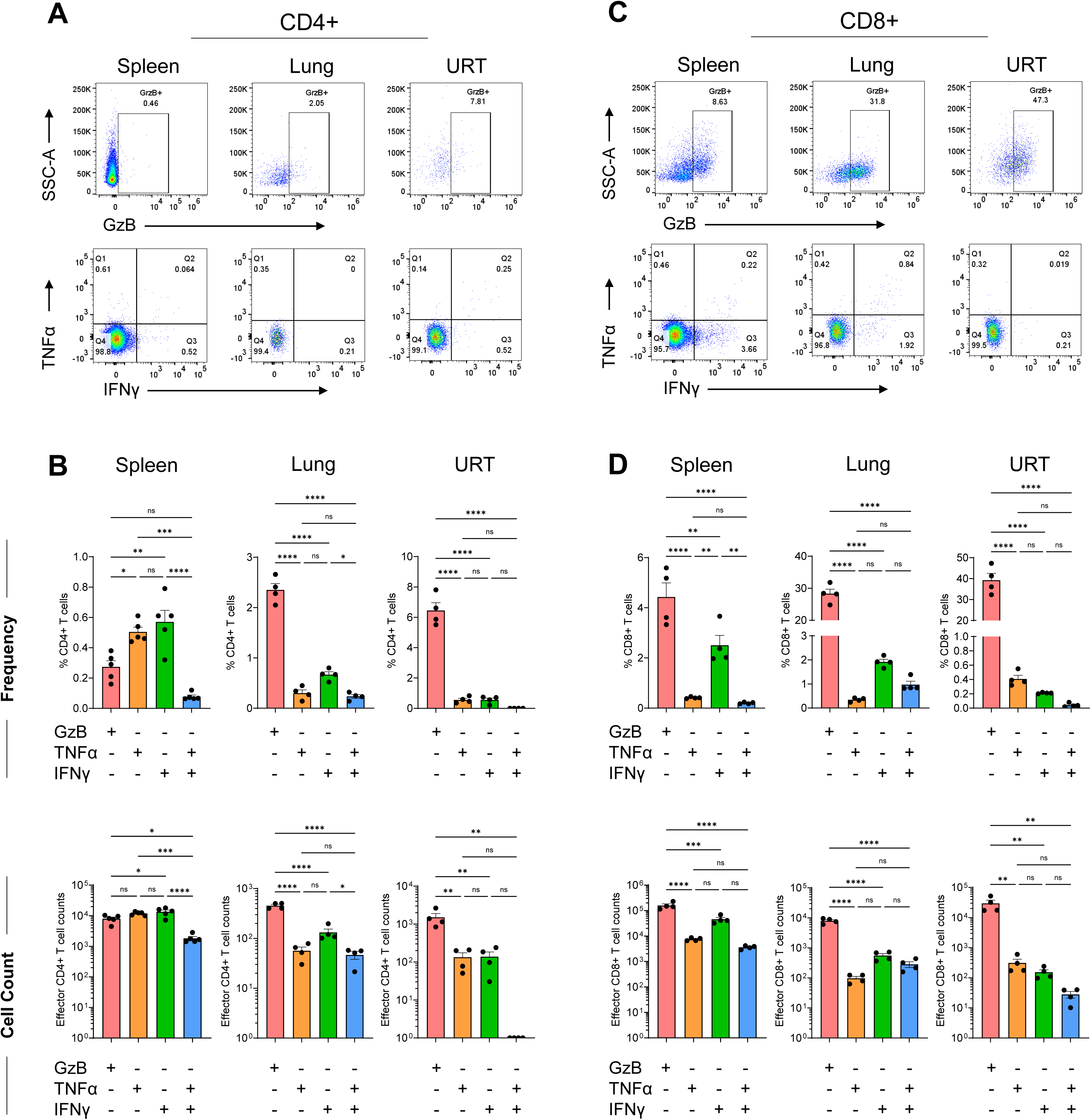
SARS-CoV-2 B.1.351 infection triggers antigen-specific T cell responses in the upper and lower respiratory tract. C57BL/6 mice were infected with the SARS-CoV-2 B.1.351 (Beta) variant at 10^6^ PFU intranasally and at day 7 pi, virus-specific CD4+ and CD8+ T cell responses were evaluated by ex-vivo peptide stimulation using spike peptide pools in spleen, lungs and URT. (A) Representative flow plots for GzB (top), TNFα and IFNγ (bottom) expression in CD4+ T cells in spleen, lungs and URT. (B) Frequency (top) and cell counts (bottom) of CD4+ T cells positive for GzB, TNFα and IFNγ expression in spleen, lungs and URT. (C) Representative flow plots for GzB (top), TNFα and IFNγ (bottom) expression in CD8+ T cells in spleen, lungs and URT. (D) Frequency (top) and cell counts (bottom) of CD4+ T cells positive for GzB, TNFα and IFNγ expression in spleen, lungs and URT. Graphs show mean ± SD. A two-way ANOVA statistical test was performed. *P < 0.05; **P < 0.01; ***P < 0.001; ****P < 0.0001; no symbol, not significant. Results are representative of data from three independent experiments with 5 mice per group.

For antigen-specific CD8+ T cells, in the lungs, nasal compartment and spleen, we observed higher cell frequencies and counts of Granzyme B secreting as compared to cytokine secreting cells (**Fig. 3c-d**). We observed 28.3% GrB+, 0.34% TNFα+, 1.91%IFNγ+, and 0.97%TNFα+IFNγ+ CD8+ T cells in the lungs, 39.25% GrB+, 0.40% TNFα+, 0.21%IFNγ+, and 0.04%TNFα+IFNγ+ CD8+ T cells in the nasal compartment, and 4.42% GrB+, 0.41% TNFα+, 2.50%IFNγ+, and 0.20%TNFα+IFNγ+ CD8+ T cells in the spleen. We compared activation markers (CD44, KLRG1, CD69) in CD4+ and CD8+ T cells across lungs, URT, and spleen (**Extended Data Fig. 2**). We discovered intriguing variations in activation between upper and lower respiratory areas. In the lungs, infected samples showed increased CD44 and CD69, and to a lesser extent, KLRG1 in contrast to mock samples. However, in the URT there was no difference in CD44 and CD69 expression in CD4+ and CD8+ T cells. Surprisingly, there was a significant rise in KLRG1 expression in CD8+ T cells in the URT. No changes were observed in expression of T cells from the spleen. These data indicate that SARS-CoV-2 infection triggers antigen-specific CD4+ and CD8+ T responses in the respiratory tract, with a higher proportion of T cells expressing Granzyme B within the respiratory tract.

### CD4+ and CD8+ T cells are dispensable for protection against SARS-CoV-2 but required for viral control within the respiratory tract

To evaluate the contribution of T cells during SARS-CoV-2 infection, we used antibody-based depletion to either individually or tandem deplete CD4+ or CD8+ T cells (**Fig. 4a**). Anti-CD4 and/or anti-CD8 antibodies were administered through the intraperitoneal route on days -5, -3, -1, +1, +7, +14, and +21 following SARS-CoV-2 infection. We assessed the efficiency of T cell depletions on days 0 and 28 pi and found near complete ablation of CD4+ and/or CD8+ T cells in whole blood, lungs and nasal compartment (**Extended Data Fig. 3a-b**). Following SARS-CoV-2 infection of CD4+ and/or CD8+ T cell-depleted mice, we observed similar peak weight loss on day 3 pi and recovery through day 10 pi. Further, we observed no mortality in any of the isotype or T cell-depleted SARS-CoV-2 infected mice (**Fig. 4b**).

**Figure 4.**
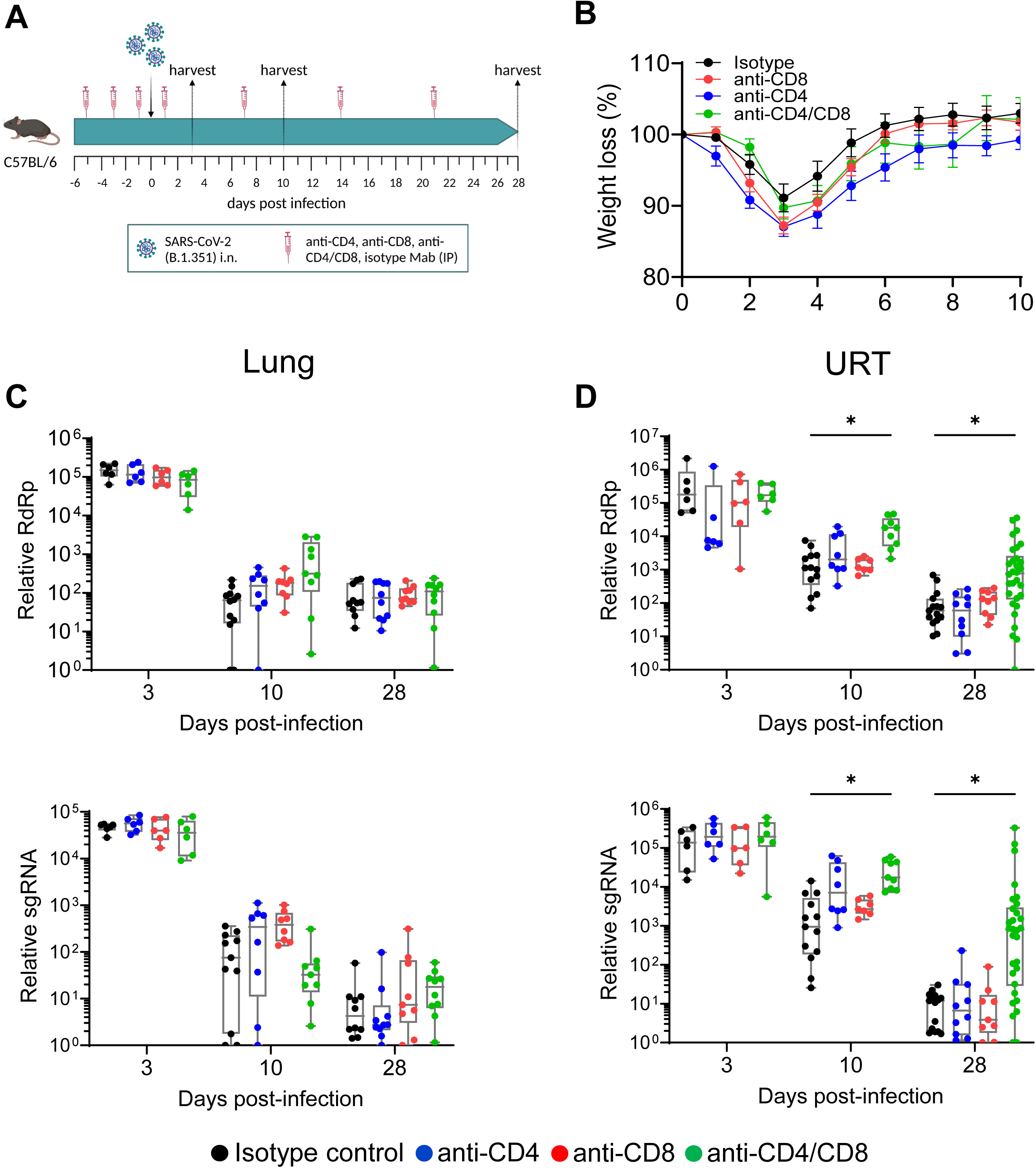
CD4+ and CD8+ T cells are dispensable for protection against SARS-CoV-2 but required for viral control within the respiratory tract. (A) Study design: C57BL/6 mice were depleted of either CD4+ or CD8+ or both T cells using 200μg anti-mouse CD4 or anti-mouse CD8α or both respectively via i.p. route at day -5, -3, -1, 1, 7, 14 and 21 post-infection. (B) Percent weight loss in SARS-CoV-2 (B.1.351) infected mice through 10 days pi Viral RNA levels as measured by relative RdRp levels (top) and sgRNA levels (bottom) at indicated time points in (C) lungs (D) URT. Graphs show mean ± SEM. Multiple Mann-Whitney statistical test was performed. *P < 0.05; **P < 0.01; ***P < 0.001; no symbol, not significant. Data are an aggregate of two independent experiments with group sizes between 6-30 mice.

We next evaluated virus replication by qRT-PCR (nasal compartment and lungs) and plaque assay (lungs). As a baseline for the depletions at later time points, we assessed viral replication on day 3 pi and observed no difference in viral RNA in the nasal compartment or lungs in the isotype or T-cell depleted mice (**Fig. 4c, d**). By day 10 pi, we observed a modest, but not statistically significant, increase in viral RNA in the lungs of the individual or tandem CD4+/CD8+ T cell-depleted mice as compared to the isotype control infected mice. In contrast, viral RNA was higher in the nasal compartment of the CD4+/CD8+ T cell tandem-depleted mice as compared to the isotype control. In the lungs by day 28 pi, the T cell depleted groups showed similar viral RNA as compared to the isotype control infected mice. However, in the nasal compartment, only the CD4+/CD8+ T cell tandem-depleted mice showed significantly higher viral RNA (30.3-fold) as compared to the isotype control infected mice (**Fig. 4c, d**).

We also compared deletion of CD8+ T cells using either CD8β or CD8α antibodies and no difference in viral RdRp levels was detected between the CD8+ T cell-depleted mice in either the CD8β or CD8α depletion group (**Extended Data Fig. 4**). These data demonstrate that T cells are required for efficient viral control/clearance within the nasal compartment, and to a lesser extent within the lungs during SARS-CoV-2 infection. Furthermore, both CD4+ and CD8+ T cells are required for preventing viral persistence within the nasal compartment and that CD4+ and CD8+ T cells can compensate for each other to control virus replication within the nasal compartment.

### SARS-CoV-2 antibody responses are CD4+ T cell-dependent but not required for viral control in the respiratory tract

Infection and vaccine-mediated antibody responses are essential protection against SARS-CoV-2 infection and have been identified as a correlate of protection^35,36^. To understand how T cells contribute to the antibody response and how these antibodies may control acute SARS-CoV-2 infection, we evaluated binding and neutralizing antibodies. On days 10 and 28 pi, we measured IgG binding antibodies to Spike, RBD and Nucleocapsid (**Fig. 5a-c**). In mice in which CD4+ T cells were depleted (individual CD4+ or CD4+/CD8+ tandem-depleted), we observed a significant reduction in anti-Spike, anti-RBD, and anti-Nucleocapsid IgG binding antibodies as compared to isotype control. In contrast, we observed no reduction in anti-Spike and anti-RBD IgG binding antibodies in the CD8+ T cell depleted mice. Furthermore, we performed a live virus neutralization assay against B.1.351 and found that CD4+ T cell depleted mice (individual CD4+ or CD4+/CD8+ tandem-depleted) showed no neutralizing antibodies above the limit of detection (FRNT_50_ = 20) (**Fig. 5d**). These findings demonstrate that the antibody response in mice during SARS-CoV-2 infection is CD4+ T cell dependent. Further, these data suggest that the antibody response likely plays little to no role in preventing viral persistence in the upper respiratory tract.

**Figure 5.**
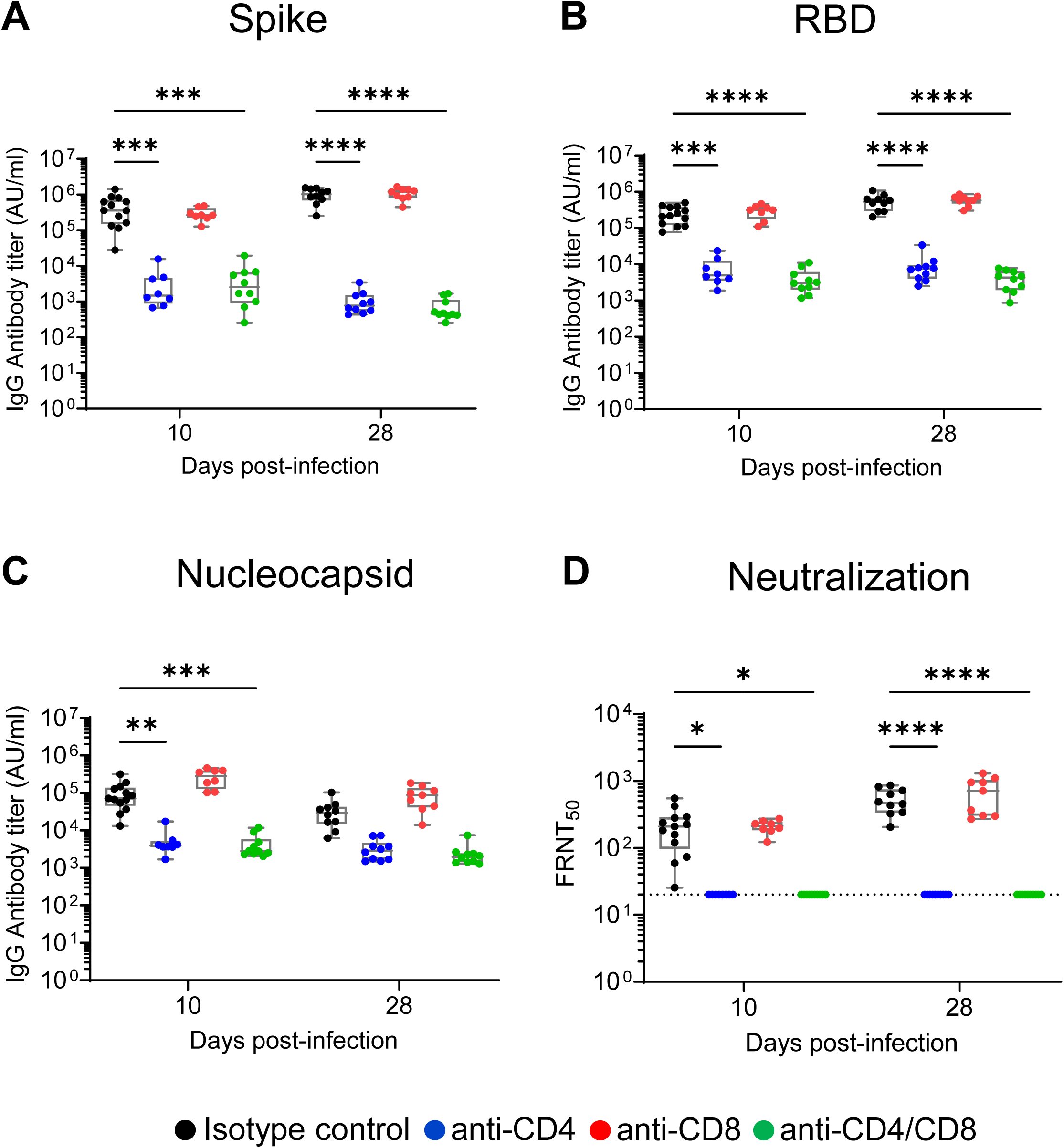
SARS-CoV-2 antibody responses are CD4+ T cell-dependent but not required for viral control in the respiratory tract. C57BL/6 mice were depleted of either CD4+ or CD8+ or both T cells and at indicated days post-infection, binding and neutralizing antibody response against SARS-CoV-2 B.1.351 spike, RBD, and nucleocapsid were measured by electrochemiluminescent multiplex immunoassay and reported as arbitrary units per ml (AU/mL) against SARS-CoV-2. IgG antibody responses were measured against (A) Spike (B) receptor-binding domain (RBD) and (C) Nucleocapsid. (D) The 50% inhibitory titer (FRNT50) on the focus reduction neutralization test (FRNT) was measured at day 10 pi and day 28 pi The dotted line in the FRNT assay represents the maximum concentrations of the serum tested (1/20). A two-way ANOVA statistical test was performedwas performed. *P < 0.05; **P < 0.01; ***P < 0.001; ****P < 0.0001; no symbol, not significant.

### SARS-CoV-2 persists within the nasal epithelium in the absence of CD4+ and CD8+ T cells

To understand whether the persistent viral RNA in the upper respiratory tract corresponds to infectious virus or not, we performed a TCID_50_ infectious virus assay from nasal compartment suspensions of isotype and CD4+/CD8+ tandem-depleted mice on day 28 pi (**Fig. 6a**). We found that 10/10 mice showed infectious virus from the tandem-depleted infected mice, but not the isotype control infected mice, with an average viral titer of 1.26 x 10^4^ TCID_50_/ml. To determine where the virus is replicating within the nasal compartment, we performed *in situ* hybridization (ISH) using RNAscope on the entire head of a mouse, which includes the nasal compartment, olfactory bulbs, and brain, with a spike-specific RNA probe (**Fig. 6a, c**). We observed no viral RNA staining within brain or olfactory bulbs in either the isotype control (n=5) or tandem depleted mice (n=10), suggesting that in the presence or absence of T cells, SARS-CoV-2 does not infect the brain. CD4+/CD8+ T cell tandem-depleted mice, but not isotype control mice, consistently showed viral RNA staining within the nasal compartment. We observed viral RNA staining within the epithelial layer and infection with little replication within the basement membrane. Additionally, infection was more commonly seen in the nasal epithelium than the olfactory epithelium found in the ethmoturbinates (**Fig. 6d**). These findings demonstrate that both CD4+ and CD8+ T cells are required to prevent viral persistence within the nasal epithelium.

**Figure 6.**
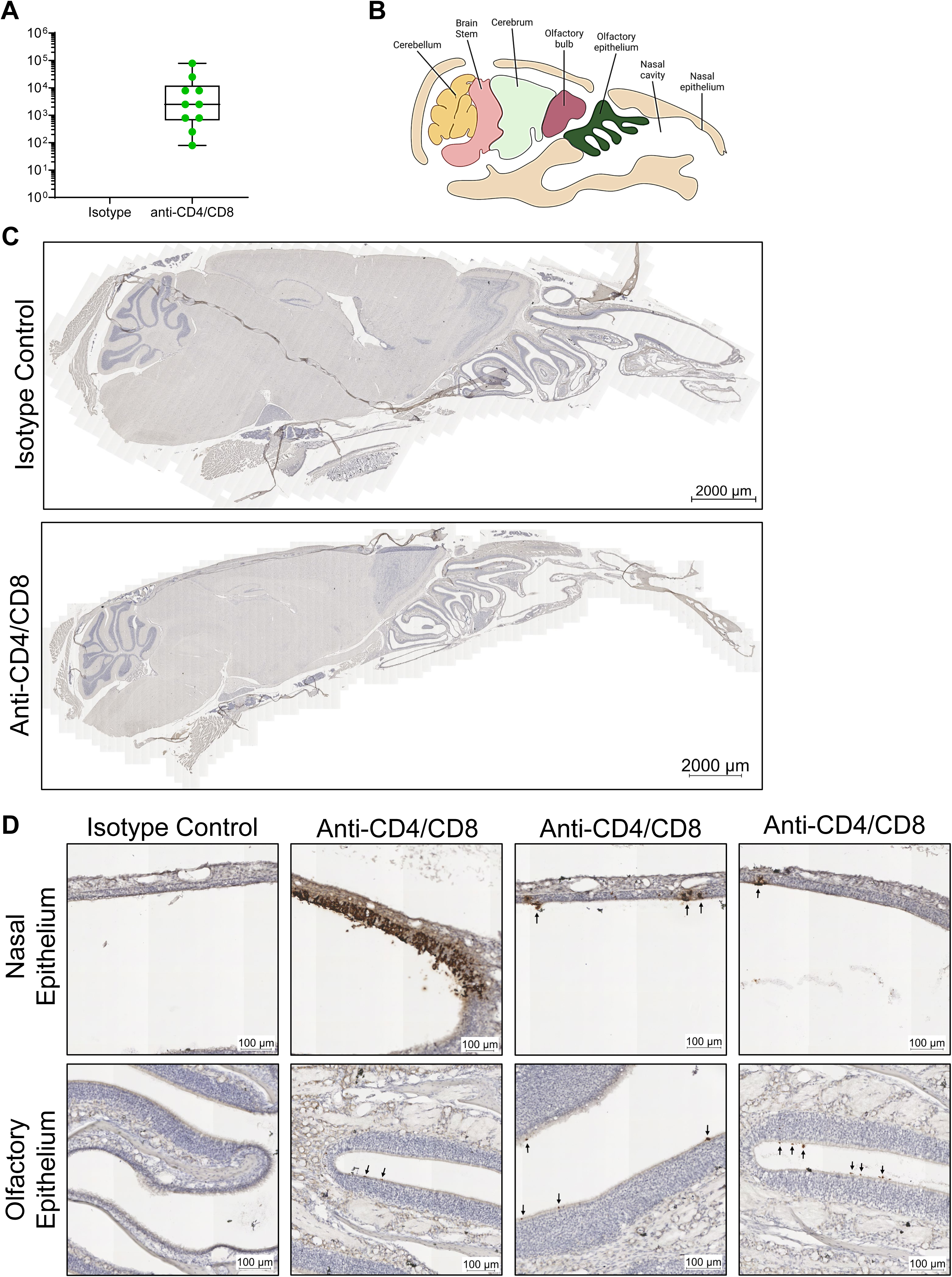
SARS-CoV-2 persists predominantly within the nasal epithelium in the absence of CD4+ and CD8+ T cells. C57BL/6 mice were depleted of both CD4+ or CD8+ and assessed for localization of viral antigen in mice heads at 28 days pi (A) Tissue culture infectious dose (TCID_50_) per mL of nasal turbinate suspension assessed in mice where both CD4+ and CD8+ T cells were depleted. (B) Representation of the sagittal section of a mouse skull showing various parts of the nasal cavity and brain (created with BioRender.com). (C) Representative images of ISH for RNA Spike Antigen performed on heads of mice where both CD4+ and CD8+ T cells were depleted and compared to Isotype control mice. (D) Representative images of nasal epithelium (top panel) and olfactory epithelium (bottom panel) of ISH for RNA Spike Antigen. Arrows represent anti-spike RNA (dark brown). Representative images from three of ten T cell-depleted mice.

### Viral persistence leads to increased viral diversity

In humans, persistent SARS-CoV-2 infection of immunocompromised individuals can lead to increased viral genetic diversity^37–41^. To understand the impact of viral persistence on SARS-CoV-2 evolution, we isolated virus on day 28 pi from the nasal turbinates of CD4+/CD8+ tandem-depleted mice and propagated it once using Vero-TMPRSS2 cells, which we have shown previously to minimize genetic diversity during virus isolation and propagation^42^. We refer to these samples as ‘isolate-28d’. Concurrently, the B.1.351 stock virus was also cultured in Vero-TMPRSS2 cells (denoted: ‘inoculum-Vero) to identify changes that may occur during a single viral passage. The ‘inoculum-Vero’ and stock virus used to infect the mice (‘inoculum-stock’) serve as the baseline for virus diversity. These sequences were compared with the stock virus used to infect the mice, referred to as ’inoculum-stock’ in the text. To account for sequence mutations resulting from mouse adaptations, mice were also infected with inoculum stock virus and samples obtained from nasal turbinates on day 3 post-infection (referred to in this text as ‘inoculum-3d) where we observe peak viral titers (Fig. 1b). These mouse isolates and inoculum controls were sequenced to characterize emergent intrahost SARS-CoV-2 variants.

To quantify the variation in the virus populations between the mouse isolates-28d and inoculum controls, we calculated the pairwise genetic distance using the L1-norm. All the three controls; the inoculum-Vero, inoculum-3d and inoculum-stock were similar in diversity to each other (**Fig. 7a**) and differed only slightly likely due to variations in their low-frequency minor variant populations (**Fig. 7a inset, Extended Data Fig. 5d-e**). Compared to the inoculum-3d controls, all virus populations in the mouse isolates significantly diverged from the inoculum-stock (p-value = 6.49e-16).

**Figure 7.**
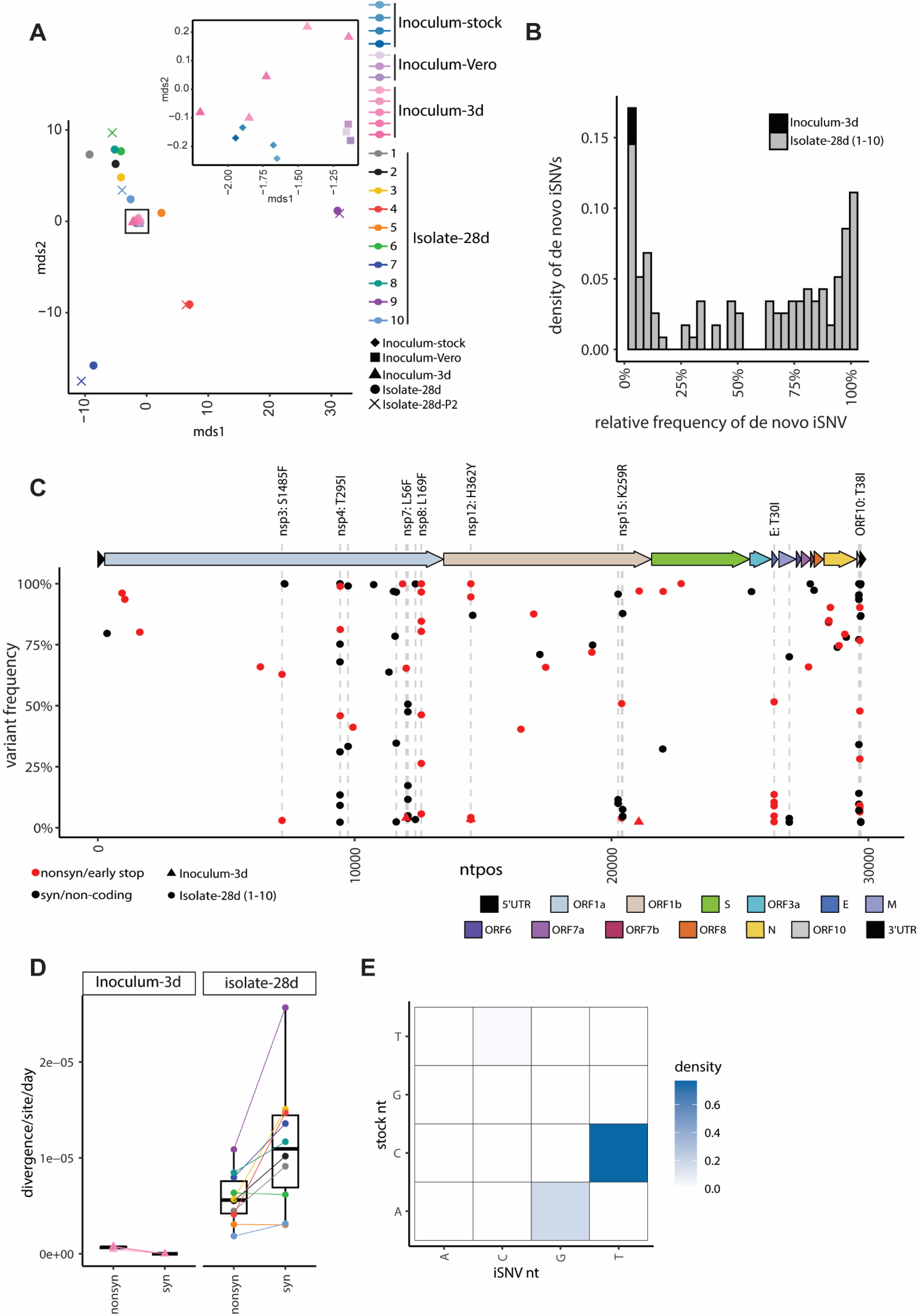
Intrahost SARS-CoV-2 variants emerge during infection in tandem CD4+/CD8+ T-cell depleted mice. (A) MDS plot of the pairwise genetic distance (L1-norm) calculations across all samples in the dataset. Point color and shape represent the sample and collection type. For the inoculum-stock, the color gradient represents different aliquots of the stock. For inoculum-Vero and inoculum-3d, the color gradients represent different infections. (B) The frequency distribution of *de novo* variants identified in the inoculum-3d (black) and isolate-28d (1-10, gray) samples. (C) The location, frequency, and characteristics of *de novo* variants in the inoculum-3d and isolate-28d (1-10) samples. The color of each point represents the type of mutation including mutations that are synonymous/in non-coding regions (black) or nonsynonymous/stop-codon mutations (red). Point shape indicates the type of sample collection. Dashed lines highlight the genomic positions where a *de novo* mutation was found in more than one mouse isolate-28d sample. Labels are added for the nonsynonymous substitutions only. (D) The nonsynonymous and synonymous divergence rates normalized by the expected number of sites in the coding sequence for the inoculum-3d samples and T-cell depleted mouse isolate-28d samples. Point color and shape represent the sample and collection type as outlined in Fig 7a. (E) The density of transitions and transversions for all *de novo* mutations (2%-100%).

To identify mutations that emerged *de novo* in the tandem T-cell depleted mice by day 28 pi, we filtered for intrahost single nucleotide variants (iSNVs) that were not present in the standing diversity of the inoculum-stock and inoculum-Vero samples (0.5%-100%). The location, number, and frequency of *de novo* iSNVs varied by mouse isolate (**Fig. 7b-c**), with isolate 10 having zero iSNVs that reached ≥ 50% while isolate 9 had 15 (6 of which were amino acid substitutions) (**Extended Data Fig. S6**). The ORF10, nsp7, and envelope had the highest mutational densities with averages of 14.5 (±5.8), 3.2 (±2.5), and 2.6 (±2.3) *de novo* iSNVs/kb, respectively. Out of the 57 unique *de novo* mutations, 19 emerged in more than one mouse, 8 of which generated amino acid substitutions in the ORF1a (nsp3, nsp4, nsp7, nsp8), ORF1b (nsp12, nsp15), E, and ORF10 gene regions. All 19 mutations were present in a small subset (<1%) of globally circulating SARS-CoV-2 sequences (CoV-Spectrum 01/06/2020-12/05/2023)^43^. Additionally, the nsp4 T295I mutation has previously been reported in mouse-adapted strains of SARS-CoV-2^17^. The remaining nonsynonymous mutations have not been characterized, though others have identified mouse-specific adaptations that emerged in nsp6, nsp7, and nsp8^44,45^. Interestingly, iSNVs shared in more than one mouse isolate were more likely to be present as minor variants (<50%) compared to iSNVs found in only a single mouse isolate (p-value = 0.0003, Fisher’s Exact), indicating that the shared iSNVs may emerge at different points during infection.

In the mice where both CD4+ and CD8+ T cells were depleted, the synonymous divergence rates (mean = 1.12e-05 per site per day) were higher than the nonsynonymous (mean = 5.84e-06 per site per day) (**Fig. 7d**, p-value = 0.043 Mann Whitney U), indicating that purifying selection is occurring in the CD4+/CD8+ tandem depleted mice. Further, the rates observed in our mouse isolates fall into similar ranges observed in immunocompromised individuals with prolonged infections (≥21 days)^46^. Over 76% of the iSNVs were C-to-T transitions (**Fig. 7e**), a signature of SARS-CoV-2 evolution observed in human intrahost studies and globally circulating strains^47^.

### Genetic diversity within SARS-CoV-2 leads to reduced virus replication in the lungs

We next selected five mice isolates based on their distance from the inoculum-stock and *de novo* iSNV populations (**Fig. 7a, Extended Data Fig. 6a, 8a**). Isolates 4, 7 and 9 diverged from the inoculum-stock the most with 13, 20, and 21 *de novo* iSNVs, respectively (**Extended Data Fig. 6**). Isolates 4 and 9 had a consensus level deletion at nucleotide positions 27264-27290 (amino acids 22-30) in ORF6 (**Extended Data Fig. 7a**). This deletion was also found at <1% in the inoculum-stock (**Extended Data Fig. 7b**). Isolate 7 had *de novo* mutations emerge in only the nucleocapsid gene (**Fig. 8a**). We propagated these five isolates on VeroE6-TMPRSS2 cells (“isolate-28d-P2”) to generate working *in vivo* stocks. After sequencing the isolate-28d-P2 samples, we confirmed that no new mutations emerged upon viral propagation except for one mutation in isolate 7 (nsp4: T461A) and two mutations in isolate 10 (nsp13: S80G, E: L37F) increased in frequency after propagation.

**Figure 8.**
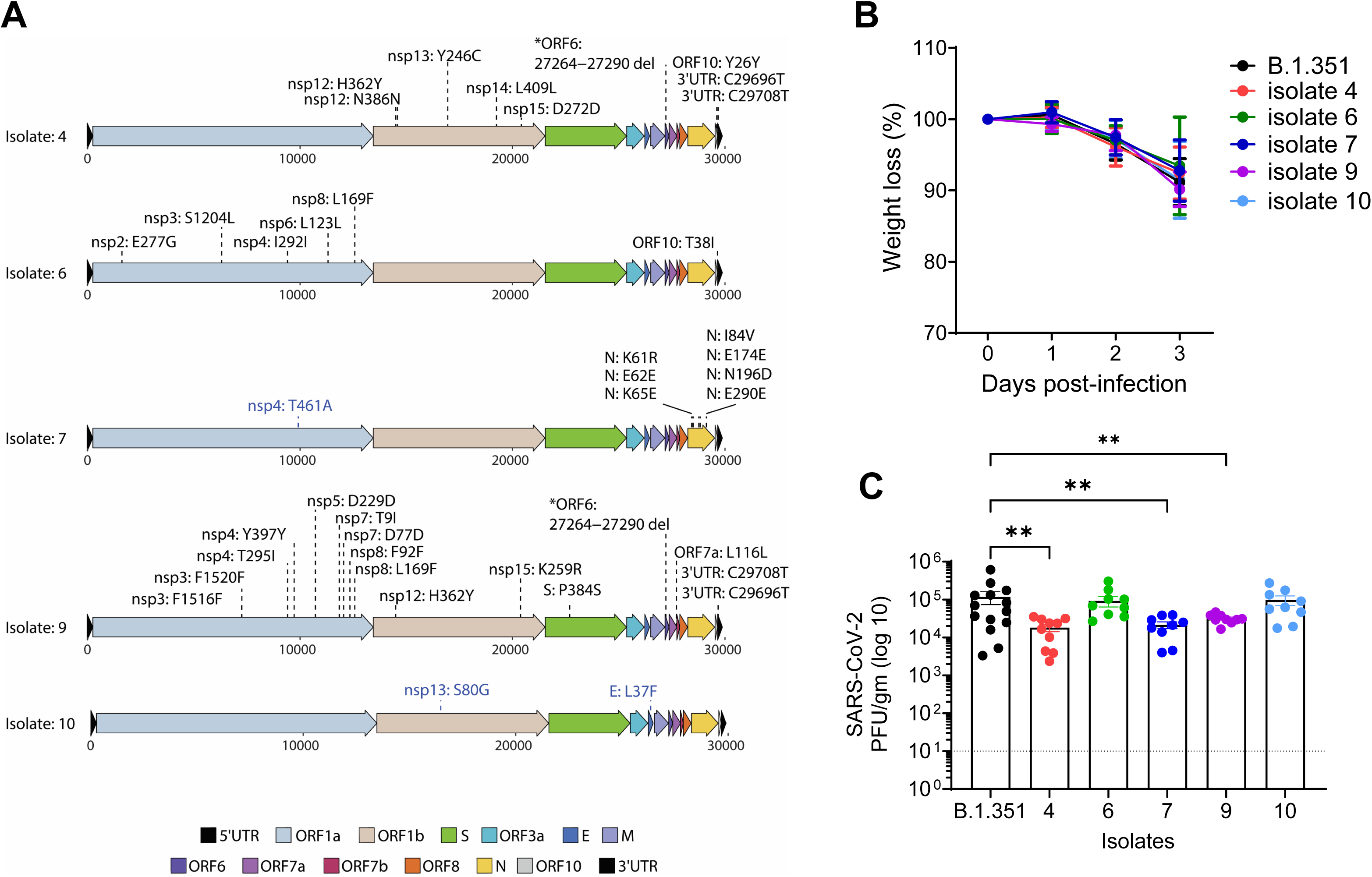
*de novo* mutations lead to differences in virus replication. (**A**) Mutation maps of *de novo* SARS-CoV-2 consensus mutations (≥50%) in the mouse isolates 4, 6, 7, 9, and 10, and compared to the isolate-Vero samples. Blue labels represent mutations that reached ≥50% in the isolate-Vero sample but were present at <50% in the original mouse isolate. Labels represent amino acid information for mutations in coding regions and nucleotide information for deletions or mutations in the non-coding regions of the genome. (B) C57BL/6 mice were infected with mice isolates-P2 as indicated and weight loss measured over three days. Graph represents percent weight loss compared to initial weight on the day of infection. (C) Infectious virus from lungs of mice infected with indicated isolates at day 3 pi was quantified by plaque assay in VeroE6-ACE2-TMPRSS2 over-expressing cells and expressed as PFU/gm of lung tissue.

We next infected C57BL/6 mice with the five mice isolates-28d-P2 at 10^6^ PFU/mL and measured weight loss throughout the infection and infectious viral titers in the lungs at day 3 pi. We observed similar weight loss dynamics across the five isolates compared to the mice infected with the B.1.351 stock virus (**Fig. 8b**). Isolate 10, which was the most similar to the stock B.1.351 virus, had similar titers to B.1.351-infected mice (**Fig. 7a, 8c**). The most divergent isolates, 4, 7, and 9, showed a 6.44, 5.44, and 3.71-fold reduction, respectively, in virus replication in the lung as compared to B.1.351-infected mice (**Fig. 7a, 8c**). Collectively, these findings demonstrate that viral persistence leads to increased SARS-CoV-2 intrahost genetic diversity and these changes can lead to differences in virus replication.

## DISCUSSION

The magnitude and quality of the T cell response is essential for driving protection against SARS-CoV-2 infection as well as promoting vaccine efficacy^48^. Several groups have investigated T cell responses against SARS-CoV-2 infection in the lungs^48–52^, however, the contribution of these cells in promoting viral control and clearance within the upper respiratory tract is not well understood. In our study, we demonstrate that T cells are necessary for controlling virus replication in the upper respiratory tract, but not the lower respiratory tract, during SARS-CoV-2 infection. Using antibodies to deplete T cells prior to infection, we found that CD4+ and CD8+ T cells play distinct roles in the upper and lower respiratory tract. Tandem depletion of both CD4+ and CD8+ T cells, but not individual, results in persistent virus replication in the nasal compartment through 28 days post-infection. The persistent virus was culturable as we were able to recover infectious virus from the nasal compartment of all tandem depleted mice nearly a month after infection. Further, we used *in situ* hybridization to determine that the virus was replicating within the nasal epithelium, and not the olfactory bulbs or brain, during persistent infection. Through deep sequencing, we found that the virus isolates from persistently infected mice show mutations across the viral genome, with several mice showing deletions within the ORF6 gene. Combined, these findings highlight the importance of T cells in controlling virus replication within the respiratory tract during SARS-CoV-2 infection.

Our studies show that T cells are critical for viral clearance within the upper respiratory tract, especially within the nasal compartment but not the lungs. In support, previous studies with the related SARS-CoV-1 virus found that B and T cells were dispensable for controlling virus replication within the lungs-highlighting the importance of innate immunity within the lungs for viral control against coronaviruses^53^. Our data also supports an important role for the innate response in controlling virus replication within the respiratory tract. Indeed, studies have found that type I and III IFNs can control SARS-CoV-2 replication in the upper and lower respiratory tracts^54,55^. Further, pDCs and alveolar macrophages have been implicated in modulating inflammation and viral control^56–59^. We recently determined the importance of the CCR2-monocyte signaling axis in promoting virus control and dissemination within the lungs and mediating protection^18^. Altogether these studies show that a combination of the innate and adaptive immune response is important for controlling virus replication, mitigating viral dissemination, limiting inflammation and protecting against lethal disease outcome.

One interesting observation was that neutralizing antibodies do not appear to play an important role in controlling SARS-CoV-2 infection. First, we show that depletion of CD4+ T cells dramatically reduces SARS-CoV-2-specific spike and nucleocapsid binding antibodies. This is consistent with studies in humans which have shown that antibody responses during SARS-CoV-2 infection and vaccination are T cell dependent^20,60,61^. Second, we show that the lack of CD8+ T cells does not alter SARS-CoV-2 spike binding or neutralizing antibody responses. Lastly, we show similar control of virus replication within the upper and lower respiratory tracts on day 10 and 28 post-infection in the absence of either CD4+ or CD8+ T cells, suggesting that antibody response is not a major driver of viral control within the respiratory tract during infection. It is possible that non-neutralizing and neutralizing antibodies play a subtle role in the kinetics of viral clearance within the respiratory tract and future studies should specifically look at the timing of the antibody response and viral control within the respiratory tract.

Under normal conditions, following respiratory virus infection, effector CD4+ and CD8+ T cells are primed in the lung-draining lymph nodes and then traffic back to the infected lung to control virus replication through their effector mechanisms^48^. There are many factors that can influence T cell programming, including but not limited to the local cytokine environment, engagement with antigen-presenting cells (APCs), interaction with co-stimulatory molecules and antigen load. Our findings suggest that T cell priming maybe different between the upper and lower respiratory tract. While Granzyme B-secreting T cells were similar between the URT and lungs, we did observe differences in cytokine-producing T cells. This suggests that anatomic compartmentalization could influence T cell responses within the respiratory tract during SARS-CoV-2 infection. Analogously, we have found during West Nile virus infection, an encephalitic RNA virus that targets the brain, shows differences in T cell activation and effector functions between the brain and spleen^62^. In the context of LCMV infection, we have previously shown that anatomic location is a driver of memory precursor CD8+ T cells into long-term memory cells within the white pulp region of the spleen^63^. Future studies should focus on identifying APC-T cell interactions within the lung-draining lymph node and within the lungs as well as how the inflammatory milieu may influence T cell responses within the upper and lower respiratory tract.

In individuals with weakened immune systems, the clearance of viruses may be delayed, resulting in prolonged shedding of the virus and accumulation of mutations in the virus genome^37,64^. One leading hypothesis is that chronic infections in immunocompromised individuals are an important source of SARS-CoV-2 genetic diversity, driving the emergence of highly divergent SARS-CoV-2 variants of concern ^37,38^. However, data are limited for studying how virus persistence, antiviral therapeutics, and host response during chronic infections impact the intrahost evolution of SARS-CoV-2. In our study, we detected viral RNA persisting in the upper respiratory tract and successfully isolated and sequenced infectious virions from the nasal turbinates 28 days post-infection. Sequence analysis of the virus populations collected from persistent infections revealed *de novo* mutations that accumulated at high frequencies across the viral genome, including mutations in ORF10 and ORF6. ORF10 had the highest density of mutations and has been shown to degrade antiviral innate immunity by mitophagic degradation of MAVS^65^. Interestingly, we observed a 27-nucleotide long deletion in ORF6 that was present at <1% in the stock inoculum but reached >50% in mouse isolates 4 and 9. The significance of the ORF-6 region in antagonizing interferon signalling during SARS-CoV-1 infections is well-established. However, conflicting findings exist regarding the role of ORF6 in SARS-CoV-2 infection, with studies showing both direct interference with IFN signaling through binding to nucleoprotein Nup98 and contradictory outcomes in animal models, including mice and hamsters^66–70^. Interestingly, there was limited diversity found in the Spike protein. Among the three observed Spike mutations, only one was present in the receptor binding domain (P384S, isolate 9) and out of the 19 mutations shared, only one (nsp4 T295I) has been identified as a mouse adaptation. Our findings, which include elevated rates of synonymous mutations and a higher occurrence of C-to-T transitions, align with observations made in prolonged SARS-CoV-2 infections of immunocompromised humans^39,47,71^. These studies collectively highlight how mutations within the viral genome can result in strategies to evade the immune system, underscoring the importance of thorough investigation into these mutations. To the best of our knowledge, our study represents one of the initial animal models illustrating SARS-CoV-2 viral persistence, leading to sequence alterations, offering a valuable tool for exploring persistent viral infections and studying the intrahost evolutionary dynamics of SARS-CoV-2.

In summary, our studies highlight the importance of T cells in mediating viral control within the respiratory tract. These findings directly impact the future design of mucosal vaccines and demonstrate the importance of promoting T cell immunity within the upper and lower respiratory tract. Further, these studies now provide a model to study innate immunity and virus-host interactions in the context of viral persistence within the upper respiratory tract.

## METHODS

### Viruses and cells

VeroE6-TMPRSS2 cells were generated as previously described^72^ and cultured in complete DMEM consisting of 1x DMEM (VWR, #45000-304), 10% FBS, 2mM L-glutamine, and 1x antibiotic in the presence of Puromycin 10mg/mL (Gibco). The SARS-CoV-2 B.1.351 variant, kindly obtained from Andy Pekosz at John Hopkins University in Baltimore, MD, was plaque purified, propagated in VeroE6-TMPRSS2 cells to create a working stock, and sequence confirmed. Viral titers were determined through plaque assays conducted on VeroE6-TMPRSS2-hACE2 cells (kindly provided by Dr. Barney Graham, Vaccine Research Center, NIH, Bethesda, MD) as described here^18^.

### Mouse experiments

C57BL/6J mice were purchased from Jackson Laboratories or bred in-house at the Emory National Primate Research Center rodent facility at Emory University. All mice used in these experiments were females between 8-12 weeks of age. Mice were anesthetized using isoflurane and infected with SARS-CoV-2 B.1.351 at 10^6^ PFU intranasally in a final volume of 50 μL in saline in accordance with the institutional standard operating procedure for working in animal biosafety level 3 facility. Post-infection, mice were monitored daily for any clinical pathology and mortality. For experiments involving T cell depletion, mice were administered 200µg/mouse of either anti-CD4 (clone GK1.5, BioXCell) or anti-CD8α (clone 2.43, BioXCell), or a combination of both anti-CD4 and anti-CD8α antibodies, or an isotype mAb IgG1 (clone HRPN, BioXCell). The administration was done via the intraperitoneal (IP) route on days -5, -3, -1, +1, +7, +14, and +21 following SARS-CoV-2 infection. Unless otherwise specified, all CD8+ T cell depletions were carried out using the anti-CD8α antibody. The impact of CD8+ T cell depletions on SARS-CoV-2 infection was also evaluated using anti-CD8β (clone 53-5.8, BioXCell) and isotype mAb IgG2b (clone LTF-2, BioXCell). All experiments and the mouse handling and care procedures followed the guidelines of the Emory University Institutional Animal Care and Use Committee.

### Antibody binding assays

Serum collected from infected mice was tested to assess the binding of IgG antibodies against B.1.351 spike, RBD, and nucleocapsid using using the V-PLEX SARS-CoV-2 Panel 7 (Mouse IgG) Kit (Meso Scale Discovery, #K15484U-2) per manufacturer protocol^72^. In brief, plates coated with the specific SARS-CoV-2 antigens were blocked using MSD blocker at room temperature, with shaking at 700 rpm for 30 minutes. Samples werediluted 1:1000 and incubated with the plates for two hours at room temperature. Following this, SULFO-TAG^TM^ conjugated Goat Anti-Mouse IgG Antibody was added. The plates were washed with 1X MSD Wash Buffer, and then MSD Gold Read Buffer B was added to each well. Between each step, the plates were washed three times with PBS containing 0.05% Tween. An MSD plate reader was used to read the plates, and the results were analyzed using Discovery Workbench® software, version 4.0 and reported as arbitrary units per ml (AU/mL) against SARS-CoV-2.

### Focus reduction neutralization assay

FRNT assays were conducted following the methods outlined in the protocol described previously^73^. In summary, the samples were diluted in 3-fold increments, creating 8 serial dilutions using DMEM in duplicates. The initial dilution was set at 1:10, and the total volume reached 60 µL. The serially diluted samples were then incubated with virus at 37°C for 1 hour in a round-bottomed 96-well culture plate. After the incubation, the antibody-virus mixture was added to Vero-TMPRSS2 cells and incubated again at 37°C for 1 hour. Following this step, the antibody-virus mixture was removed, and a 0.85% methylcellulose overlay (Sigma-Aldrich) was added to each well. The plates were further incubated at 37°C for 18 hours. Once the incubation was complete, the methylcellulose overlay was removed, cells were washed with PBS and fixed using 2% paraformaldehyde in PBS. Plates were washed with PBS and permeabilized using a buffer containing 0.1% BSA and Saponin in PBS for 20 minutes. Following this, cells were incubated overnight at 4°C with the anti-SARS-CoV spike primary antibody directly conjugated to Alexa Fluor-647 (CR3022-AF647). Subsequently, the cells underwent two washes with 1x PBS before imaging on an ELISPOT reader (CTL Analyzer).

### TCID50 assay

VeroE6-TMPRSS2 cells were seeded at a density of 25,000 cells per well in complete DMEM media in quadruplicates for each sample. Once confluent, the medium was removed, and 180 µL of DMEM containing 2% FBS and was added. Serial dilutions of samples, along with positive controls (virus stock with a known infectious titer) and negative controls (medium only), were included. The plates were further incubated at 37°C with 5.0% CO2 for 2 to 5 days. Cells were fixed and stained with a crystal violet solution containing 2% paraformaldehyde. Visual inspections were carried out on cell monolayers to detect any cytopathic effect and TCID50 was determined using the Read–Muench formula^74^.

### *In situ* hybridization of brain tissues

Mice heads collected on day 28 pi were subjected to decalcification in ethylenediaminetetraacetic acid (EDTA) for approximately two weeks, followed by rehydration in PBS for two days. Subsequently, the samples were immersed in a 30% sucrose solution prepared in PBS for three to four days to achieve sucrose equilibration. Once equilibrated, the heads were rapidly frozen in 100% Optimal Cutting Temperature (O.C.T.) compound and stored at -80°C. Tissues were sliced using a cryostat and stored at -80°C. RNA in situ hybridization (RNA-ISH) was performed following the RNAscope Brown Kit protocol, with a modification for OCT-frozen tissues where tissue is post-fixed onto the slide. The tissue was stained for Spike RNA using a custom RNAscope Probe V-nCoV2019-S (ACD, #848561). Images were obtained using a Zeiss AxioImager Z2 system with Zeiss software at 20X on the Slide Scanner, an automated imaging system.

### Processing of mouse tissues for flow cytometry analysis

On the specified day following infection, mice were anesthetized using isoflurane and given a retro-orbital injection of CD45:PE (BD Biosciences, clone 30-F11) diluted 1:20 in saline in a final volume of 100 μL per mouse. After a 5-minute recovery period, the mice were euthanized using isoflurane overdose. Lung, URT tissues and spleens were harvested from each mouse and placed in 1% FBS-HBSS. The spleens underwent mechanical homogenization on a 70 μM cell strainer, and the resulting cell suspension was collected in 10% FBS-RPMI. Splenocytes were processed by centrifugation (1250 rpm, 5 min, 4C), followed by lysis in ACK Lysis buffer for 5 minutes on ice. After washing with 10% FBS-RPMI, the splenocytes were kept chilled until they were ready for further applications. The lungs were mechanically disrupted in 6-well plates and then subjected to a 30-minute digestion at 37C using a solution of DNaseI and collagenase in HBSS. The digestion was halted with 10% FBS-RPMI, and the lung cells were filtered through a 70 μM filter to obtain a single-cell suspension. The obtained cells were layered onto a Percoll-PBS gradient, centrifuged, and the top layer of cell debris was removed. The resulting cell pellet was lysed with ACK lysis buffer for 5 minutes on ice, followed by washing and resuspension in 10% FBS-RPMI, keeping the cells chilled until they were ready for the staining process.

### Flow cytometry analysis

Single-cell suspensions were centrifuged and were then resuspended in a blocking solution containing anti-CD16/32 (Tonbo, Clone 2.4G2) for 20 minutes at 4°C. After centrifugation, the cell suspensions were stained using surface stain antibodies including Live/Dead Ghost Dye stain (Tonbo Biosciences) for 20 minutes at 4°C. Following this, the stained cells were washed and fixed in 2% PFA-PBS for 30 minutes at room temperature. Finally, cells were washed, resuspended in 300 μL of FACS buffer (1% FBS in 1x PBS). Precision count beads (Biolegend) were added to the samples to facilitate cell counting. These samples were then processed using a BD FACS Symphony A5 instrument. The anti-mouse surface staining antibodies utilized in this study were: CD45:PE (Biolegend, Clone: 30-F11), CD45.2:BV650 (Biolegend, Clone: 104), CD44: FITC (Biolegend, Clone: NIM-R8), CD3e: PerCP Cy5.5 (Tonbo, Clone: 145-2C11), IL-7R: PE/ Cyanine 5 (Biolegend, Clone: A7R34), KLRG1:APC-Cy7 (Biolegend, Clone: 2F1), CD8b: BV421 (Biolegend, Clone: YTS156.7.7), Live/Dead Ghost Dye™ UV 450 (Tonbo), CD69: BV785 (Biolegend, Clone: H1.2F3), CD103: AF700 (Biolegend, Clone: 2E7), CD4: PE-Cy7 (Biolegend, Clone: GK1.5).

### *Ex vivo* T cell assays

For T cell peptide restimulation, approximately 1x10^6^ cells harvested from tissues were placed per well in a 96-well round bottom plate and stimulated for six hours at 5% CO2, 37°C, with the addition of 10 μg/mL brefeldin A (BFA) in complete RPMI media. For positive control, splenocytes were stimulated with BFA and PMA/Ionomycin. For negative control, cells were stimulated with BFA and vehicle DMSO in the same media. To measure antigen-specific T cell responses, splenocytes, URT and lung cells were stimulated with 1 μg/ml of SARS-CoV-2 spike peptide pool (BEI resources) in the presence of BFA. After T cell stimulation, cells were washed with FACS buffer and stained for the surface antigens as described in the previous section. Cells were then washed with FACS buffer and incubated with 1x Fix/Perm solution (Tonbo) at room temperature for one hour. Following this, cells were washed with 1x Perm buffer (Tonbo) and stained for intracellular antigens for 30 mins at 4°C with the following antibodies: GranzymeB: AF647 (Biolegend, Clone: GB11), IL-2: BV605 (Biolegend, Clone: JES6-5H4), TNFα (Biolegend, Clone: MP6-XT22) and IFNγ: PE Dazzle 594 (Biolegend, Clone: XMG1.2). Following intracellular staining, cells were washed twice with the 1x Perm buffer and once with FACS buffer. Cells were resuspended in 300 μL of FACS buffer. Precision count beads (Biolegend) were added to the samples to facilitate cell counting. These samples were then processed using a BD FACS Symphony A5 instrument. For data analysis, splenocytes were gated on lymphocytes, single cells, live, parenchymal lymphocytes, CD3+ and then further categorized as either CD4-, CD8+ for CD8 T cell response analysis, or CD4+, CD8-for CD4 T cell analysis. Antigen-specific cells were identified based on their production of IFN-γ, TNF-α, or both cytokines in response to SARS-CoV-2 peptide restimulation.

### Quantitative reverse transcription-PCR (qRT-PCR)

To prepare tissue samples for evaluating viral RNA levels and mRNA expression, lung and nasal turbinate tissues were collected in an Omni Bead Ruptor tube containing Tri reagent (Zymo). Subsequently, tissues were homogenized using the Omni Bead Ruptor 24 instrument with a program set at 5.15 m/s for 15 seconds. The samples were then stored at -80°C until further analysis. Samples in Tri reagent were briefly spun and then RNA extracted using Direct-zol RNA miniprep kit (Zymo) and cDNA was prepared using High-Capacity cDNA Reverse Transcription Kit (Thermo Fisher Scientific) as per the manufacturer’s protocol. Viral RNA levels and replication were measured as previously described^54^. Briefly, qRT-PCR was set up using IDT Prime Time gene expression master mix on a QuantStudio5 qPCR system using the cycling conditions recommended by the manufacturer. To measure viral RNA levels, SARS-CoV-2 RDRP-specific forward primer: GTGARATGGTCATGTGTGGCGG; reverse primer: CARATGTTAAASACACTATTAGCATA, and probe 56-6-carboxyfluorescein [FAM]/CAGGTGGAA/ZEN/CCTCATCAGGAGATGC/3IABkFQ were used. To measure virus replication, levels of SARS-CoV-2 E gene subgenomic RNA (sgRNA) was measured using forward primer sgLeadSARSCoV2-F: 5’-CGATCTCTTGTAGATCTGTTCTC-3’ (IDT) and the E_Sarbeco R2 reverse primer (IDT; #10006890) and P1 FAM probe (IDT; #10006892). GAPDH was used as a reference gene to normalize viral RNA levels which were represented as fold change over mock samples.

### Illumina library preparation, sequencing, and alignment

SARS-CoV-2 RNA was isolated using RNAzol® RT Column Kit (Molecular Research Center, Inc.) as per manufacturer’s instructions from the B.1.351 stock samples (n=4, “inoculum-stock”), B.1.351 stock samples passaged once in VeroE6-TMPRSS2 cells (n=3, “inoculum-Vero”), nasal turbinates of C57BL/6 control mice at day 3 pi (n= 5, “inoculum-3d”), nasal turbinates of CD4+/CD8+ tandem depleted mice on day 28 pi and passaged once in VeroE6-TMPRSS2 cells (n=10, “isolate-28d”), and five mouse isolates passaged once more in VeroE6-TMPRSS2 cells (“isolate-28d-P2”). Approximately 400bp long amplicons were generated from the isolated SARS-CoV-2 viral RNA using the ARTIC V4 primers and protocol (https://artic.network/2-protocols.html). Amplicons were cleaned using AMPure beads and input into the Illumina DNA Prep Kit (Illumina, San Diego, CA) according to the manufacturer’s protocol. The concentration and fragment size of the libraries were determined using the Qubit dsDNA high-sensitivity assay (ThermoFisher Scientific, Waltham, MA) and a high-sensitivity D1000 screentape (Agilent, Santa Clara, CA), respectively. The final libraries were pooled at equal molarity and sequenced on the MiSeq (v3 600 cycles, Illumina, San Diego, CA). Amplification, library preparation, and sequencing were done twice on the same RNA sample.

Reads were trimmed with trimmomatic v0.39^75^and aligned to the Wuhan/Hu-1 SARS-CoV2 genome (NC_045512.2) using bwa mem v0.7.17^76^. ARTIC v4 primer sequences were removed using iVar v1.3.1^77^ with a minimum quality threshold of 0, including all reads with no primer sequences found. Consensus sequences and variants were called using an in-house variant calling pipeline, timo (v4) (https://github.com/GhedinLab/timo)^78^. The amplification and library preparation protocol and alignment pipeline are available at https://github.com/GhedinSGS/SARS-CoV-2_analysis. The sequencing data are available at (PRJNA1064978).

### Single nucleotide variant analysis

Coverage and variant data were pulled from the timo outputs (**Extended Data Fig. 5a-c**). Minor variants in the inoculum and inoculum-Vero samples were required to be present in both sequencing replicates with an average frequency of 0.5%-49% and read depth of 200X. Nucleotides ≥50% at positions with ≥ 10X read depths (dp) were considered the consensus nucleotide for the sample. All consensus sequences for the inoculum-stock and inoculum-Vero samples were confirmed to be identical. Passaging the inoculum stock once in the VeroE6-TMPRSS2 filtered out a large proportion of low-frequency (mean = 1%, median = 0.7%) minor variants in the inoculum-stock used for infections (**Extended Data Fig. 5d-e**). Therefore, intrahost single nucleotide variants (iSNVs) that were generated *de novo* were required to be present at 2%-49% and at positions with 200X dp for minor variants or 50%-100% 10X dp for consensus variants, and not present in any inoculum stock or inoculum-Vero samples, including as a minor variant at 0.5%-49%. Tables outlining the *de novo* iSNVs are located at https://github.com/GhedinSGS/TCD_Mice.

### Genetic distance and divergence calculations

Divergence rates were calculated as previously outlined^79^. The number of nonsynonymous and synonymous sites were estimated for each sample using the number of nucleotide positions in the coding sequence with at least 200X read depth. All mouse isolates had >90% coverage of the coding sequence at 200X. Sites that lacked any minor variant and were identical in their consensus nucleotide across all samples were not used for distance calculations, as the distance for these sites equals 0. All other sites were used to calculate the pairwise genetic distance using the L1-norm. Data and code are located at https://github.com/GhedinSGS/TCD_Mice.

### Quantification and Statistical Analysis

All experiments in mice were repeated at least twice, with representative results from one experiment shown. Statistical analysis was performed in GraphPad Prism 10.1.1 (Prism, La Jolla, CA, USA) using Student’s t test to compare two groups and unpaired one-way analysis of variance (ANOVA) to compare more than two groups. Statistical significance was defined as *P* values less than 0.05.

Antibody neutralization was quantified by determining the foci count for each sample done in duplicates with the aid of the Viridot program^80^. To calculate the neutralization titers, the following formula was applied: 1 - (mean number of foci in the presence of sera divided by foci at the highest dilution of the corresponding sera sample). The FRNT-50 titers were estimated through 4-parameter nonlinear regression in GraphPad Prism 8.4.3. Samples that did not exhibit neutralization at 50% were plotted at 20 and used for calculating the geometric mean.

RT-qPCR results are expressed relative to mouse Gapdh expression for the same sample and were calculated using the ΔΔCT relative quantitation method as compared to mock age-matched controls.

## Acknowledgements

This work was supported in part by grants (P51 OD011132 to Emory University) from the National Institutes of Health (NIH), Emory Executive Vice President for Health Affairs Synergy Fund award, the I3 Synergy Awards, COVID-19 cures, CEIRR HHSN272201400004C and CEIRR 75N93021C00017. This work was also supported in part by the Division of Intramural Research of NIAID/NIH.

## Author Contributions

M.K. and M.S.S. contributed to the acquisition, analysis, and interpretation of the data, as well as the conception and design of the work, and writing of the manuscript. K.E.E.J. contributed to the acquisition, analysis, and interpretation of the data, and writing of the manuscript. K.F. and A.P. contributed to the acquisition, analysis, and interpretation of the data. A.V., E.G, E.T.S., E.S., S.B, W.W., S. Sathish, S. Shrihari, and M.D.G., contributed to the acquisition and analysis of the data. A.P., R.K., A.G. and E.G., contributed to the interpretation of the data and conception and design of the work.

## Notes

### Competing Interest Statement

The authors have declared no competing interest.

