## Supplementary figures and images for "CD4+ and CD8+ T cells are required to prevent SARS-CoV-2 persistence in the nasal compartment"

### Extended Figures

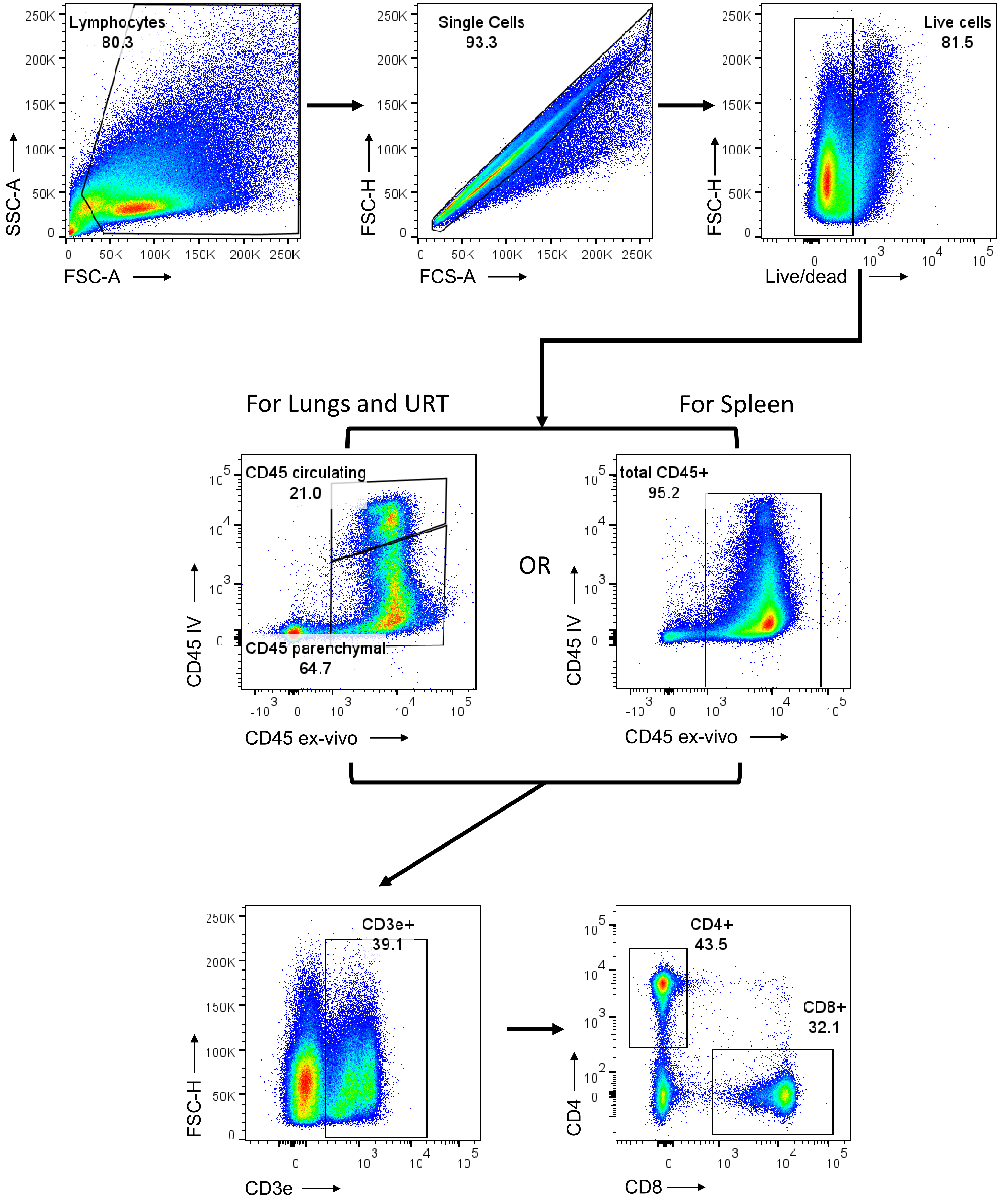

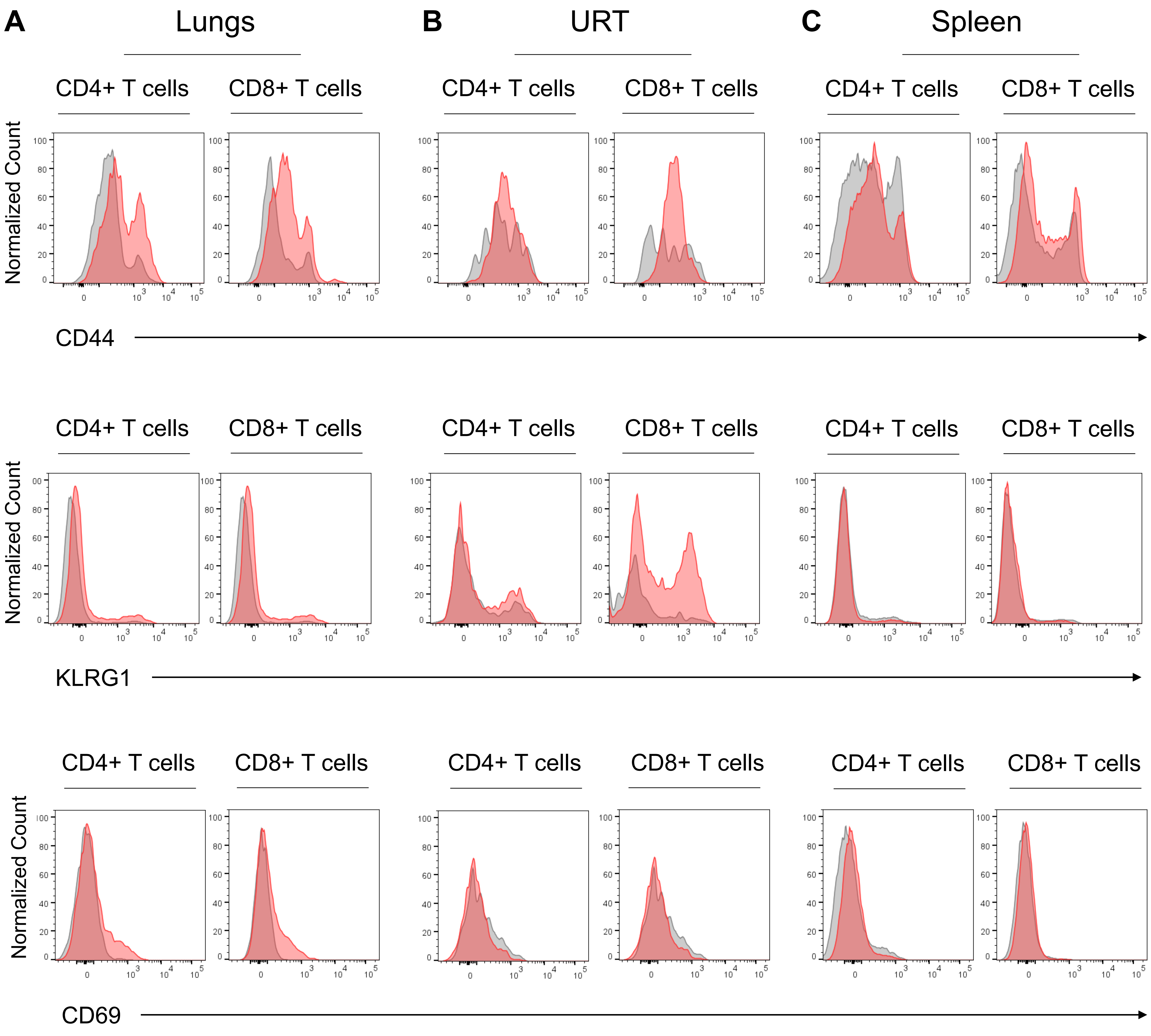

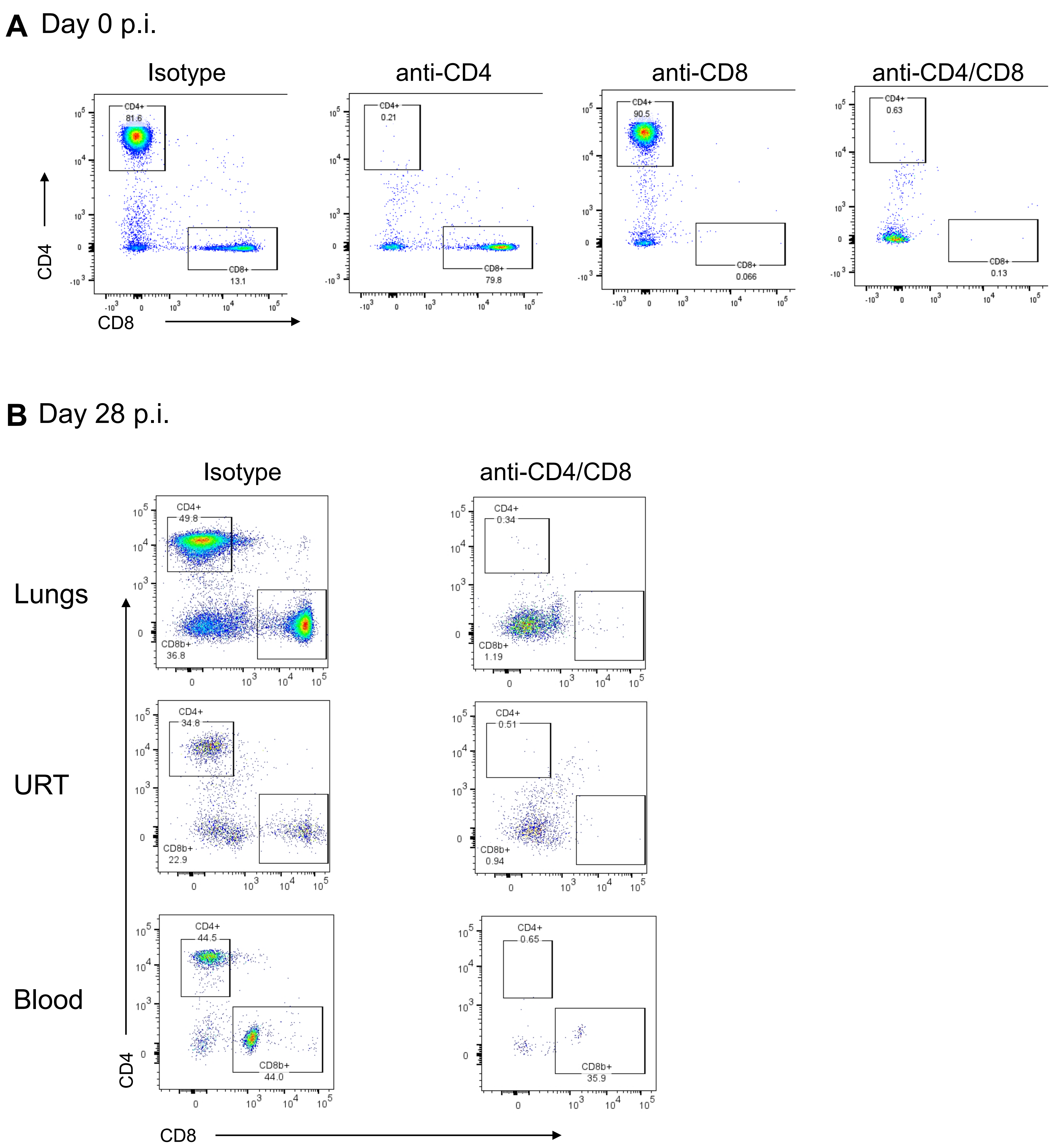

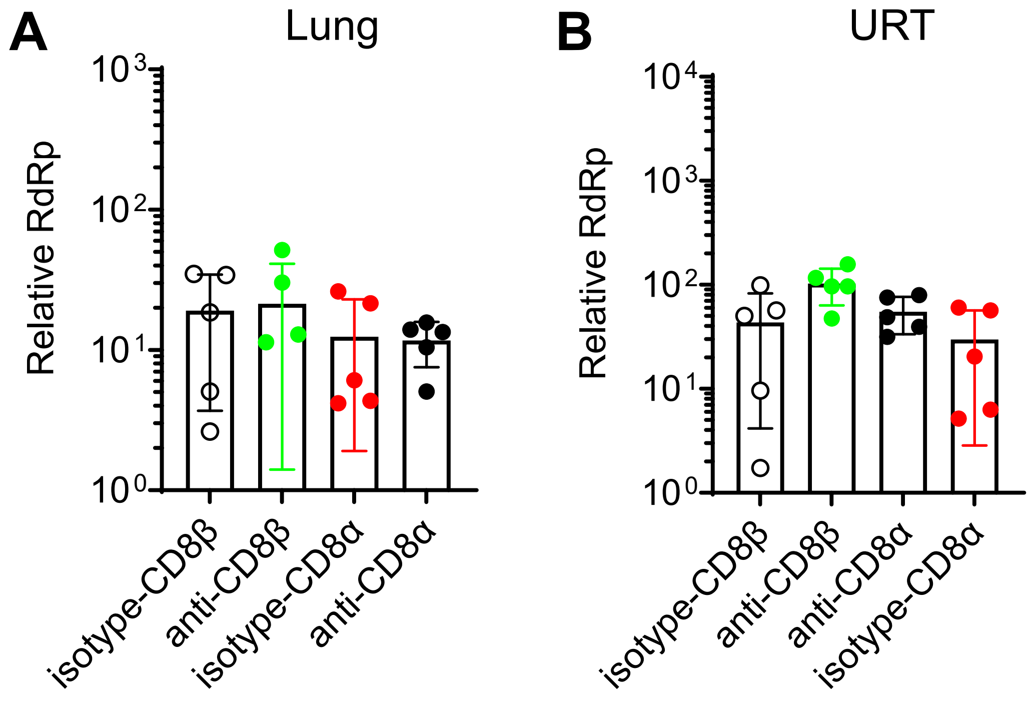

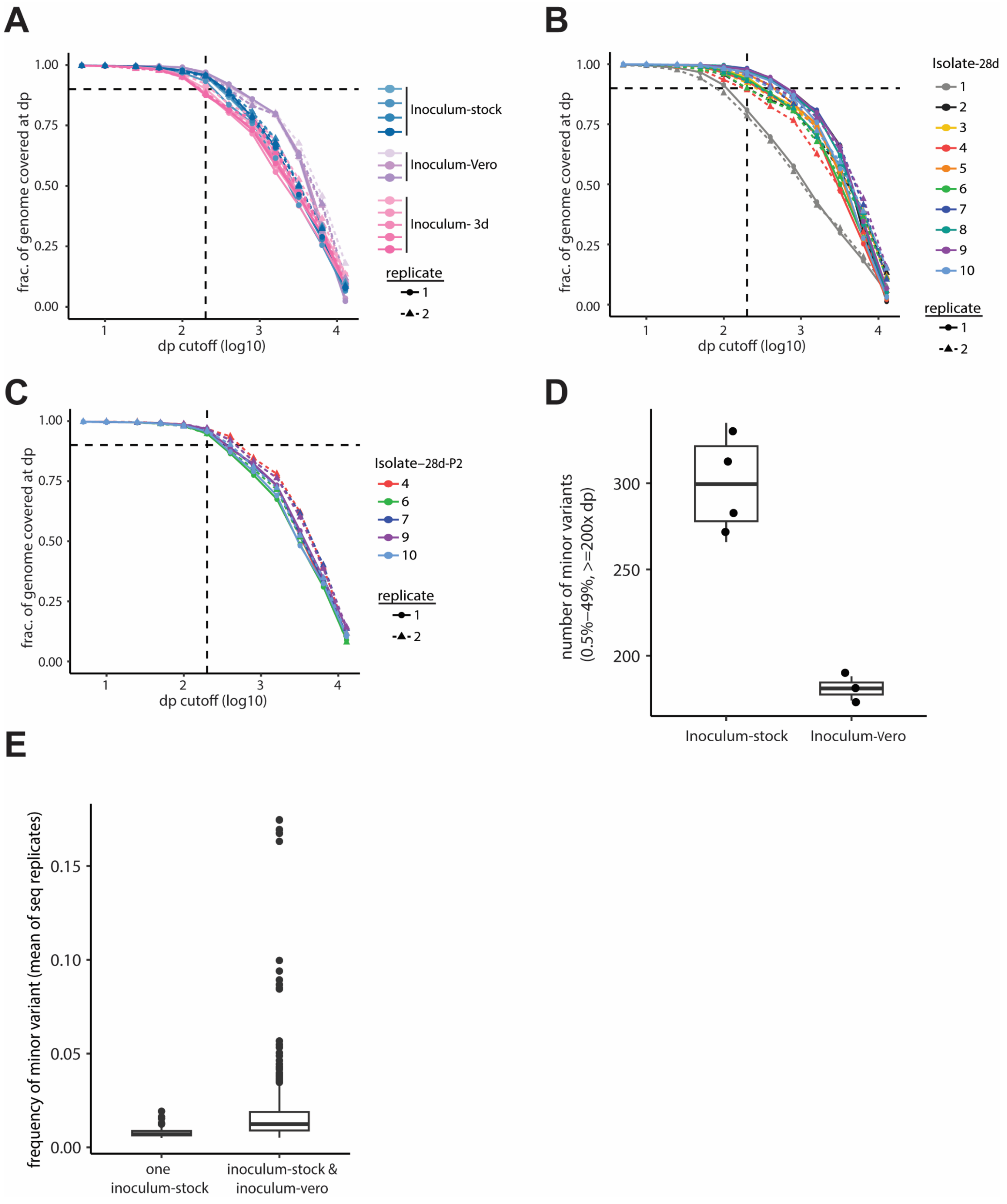

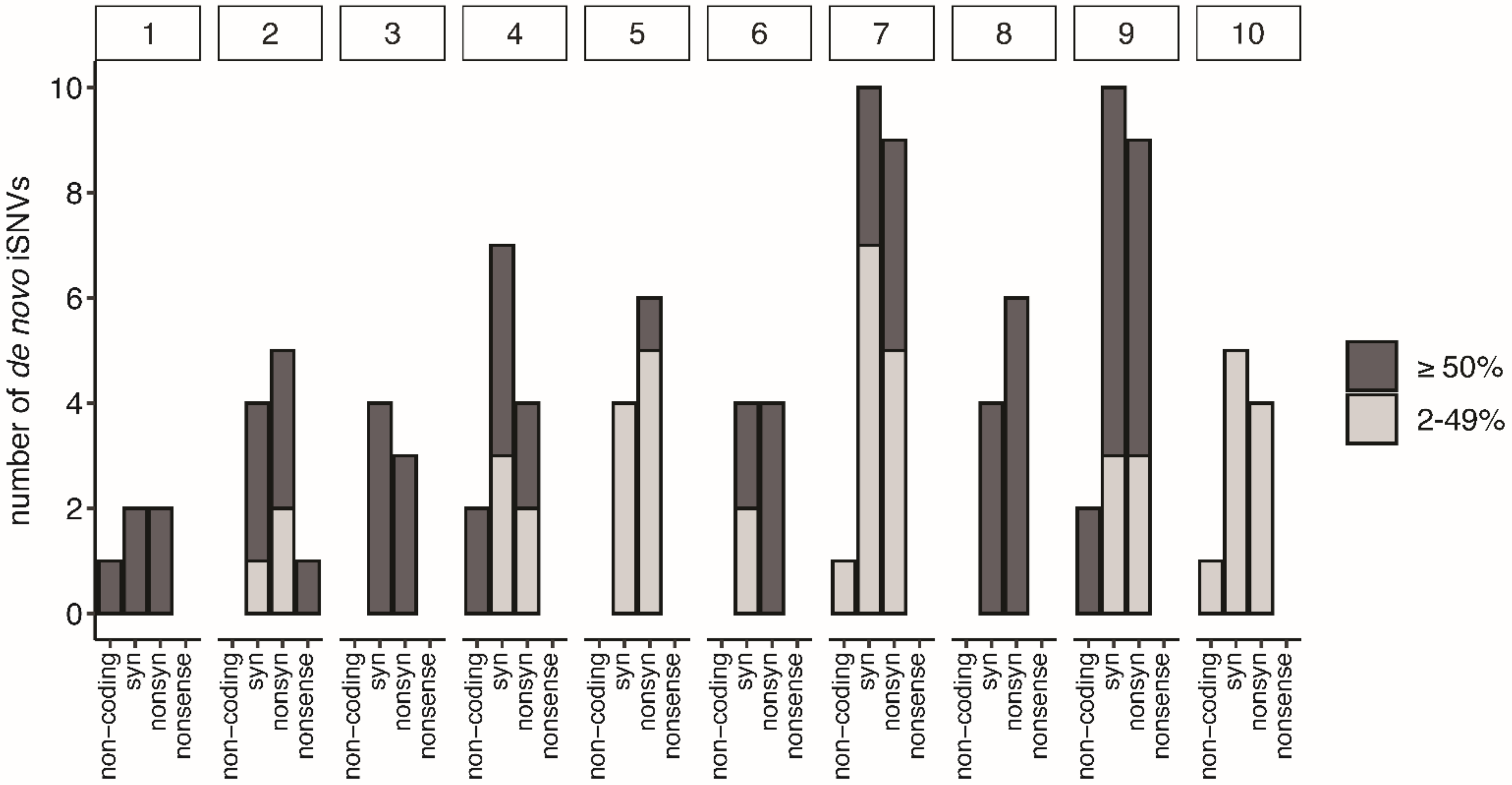

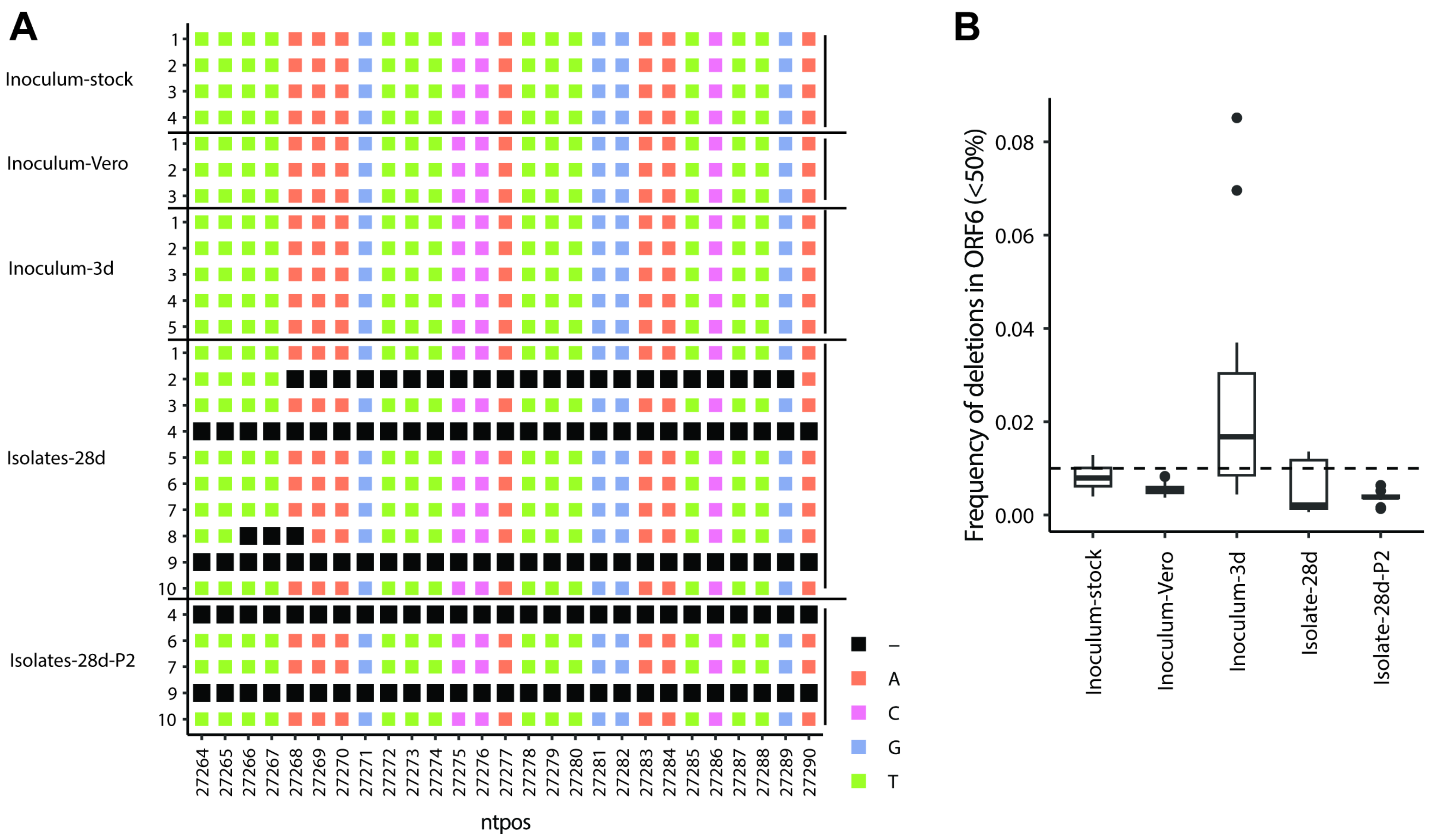
